# Genetics and genomics of enteroarthroconidial development in *Coccidioides*

**DOI:** 10.64898/2026.10.27.762035

**Authors:** Kristopher E. Fleming, Kaylee C. Christensen, Edyta Szewczyk, Christina M. Homer, Rachel B. Brem

## Abstract

Regulated cell death is a cornerstone of plant and animal development. Distant orthologs of some cell death genes are also found in microbes, with roles that remain incompletely understood. The human fungal pathogen *Coccidioides* causes the disease Valley fever when infectious spores called enteroarthroconidia are inhaled from soil. These spores develop via an unusual checkerboard-style differentiation in which alternating segments of hyphae (cellular filaments) die and disarticulate, releasing mature spores on either side. The molecular mechanisms are unknown. We first adapted single-cell transcriptomics for filamentous fungi, cataloguing expression across the specialized cells of a developing *Coccidioides* colony. With the resulting profiles, and cell-biological validation via protein tagging, we characterized high expression of the pyroptosis-like cell death pathway in hyphal segments destined to die. Next, we tested the function of pyroptosis-like pathway genes directly by genetic knockout; strains lacking these factors exhibited a phenotype of developmental arrest at a late stage in the colony, preserving hyphal growth but abrogating cell death and spore production. Together, these findings reveal the pyroptosis-like pathway as a requirement for enteroarthroconidial development, and thus infectivity, in *Coccidioides.* We propose that regulated cell death may prove to mediate events in development in many fungi with multicellular life cycles.

**Author summary:** Regulated cell death is a cornerstone of development in plants and animals. Some of the underlying genes have distant orthologs in microbes, with functions that remain incompletely understood. In this work, we report that the pyroptosis-like cell death pathway is required for production of infectious spores by the fungus *Coccidioides*, which causes the human disease Valley fever. This first-ever molecular dissection of *Coccidioides’* infectivity has applications in Valley fever prediction and prevention. And our findings have broader biological impact as an anchor for a principle in which regulated cell death plays key developmental roles in fungi.

## Introduction

The fungus *Coccidioides* is the causal agent of Valley fever, a treatment-refractory and sometimes deadly disease prevalent in arid regions of the Western Hemisphere. Patients contract Valley fever by inhaling *Coccidioides* spores, called enteroarthroconidia, from desert soil. Valley fever incidence has risen sharply in recent years, for reasons that are poorly understood [1]. Understanding mechanisms of *Coccidioides’* infectivity is a linchpin of public health measures for disease prediction and prevention.

Enteroarthroconidia are a very rare morphology in the fungal kingdom, found only in *Coccidioides* and its close relatives. They develop in aerial hyphae, interconnected cellular filaments that grow upwards from the growth substrate. Alternating hyphal segments convert into spores, and the flanking cells, named disjunctor cells, are emptied of their cytoplasm, die, and disarticulate [2] (Fig 1a). This cell death event has remained a mystery in the field: against the backdrop of classic studies of other arthroconidial types [3,4], the molecular basis of the enteroarthroconidial strategy is unknown. Decades of work have characterized the biology of regulated cell death in plants and animals, particularly in developmental and immune processes [5]. Distant orthologs of regulated cell death genes have also emerged in fungi, implicated in allorecognition, immunity, and kin protection [6–8] and, in the plant pathogen *Magnaporthe oryzae*, in infection initiation by fungal spores [9,10]. Developmental roles in multicellular fungal lifecycles, like that of *Coccidioides*, have been uncharacterized.

**Fig 1.**
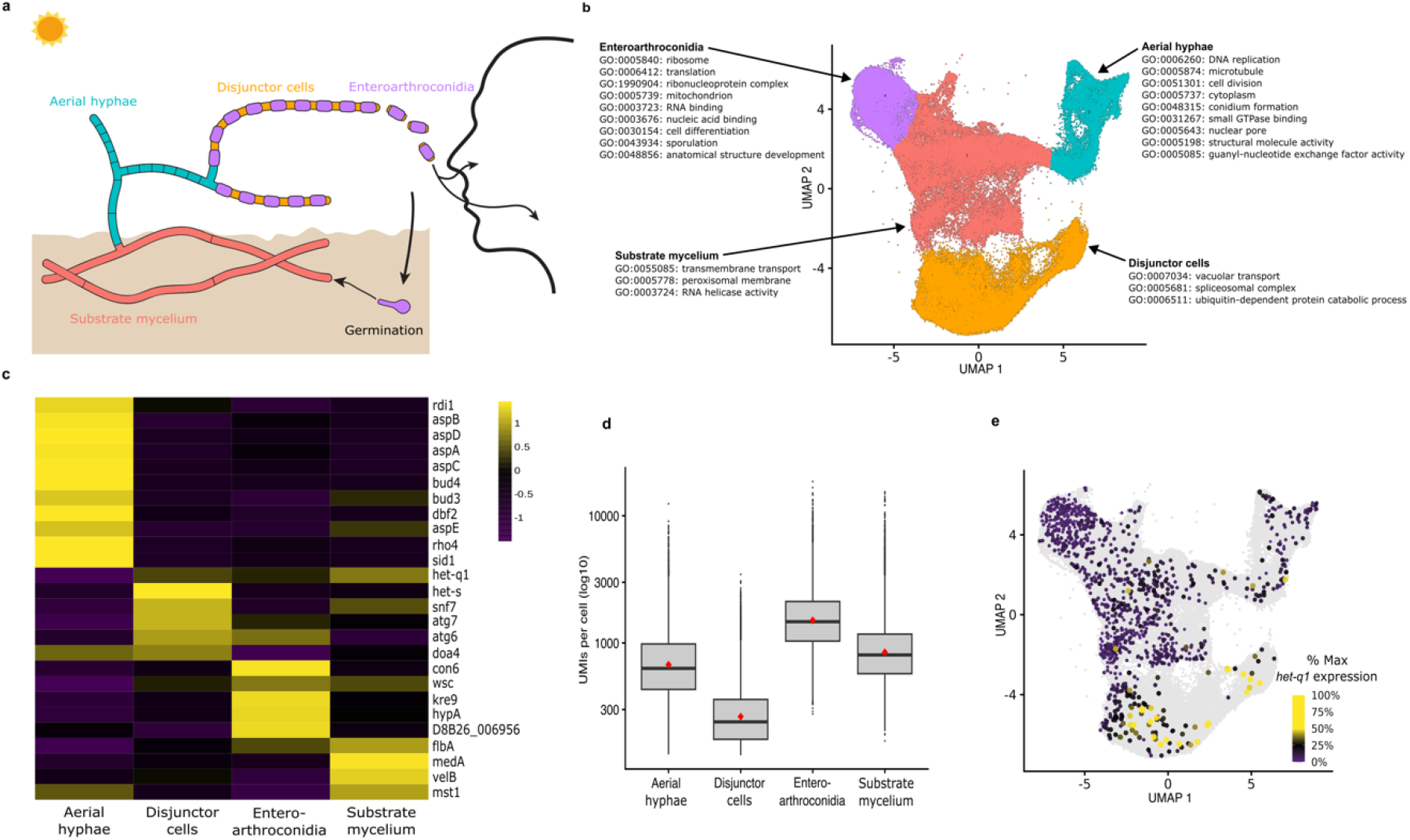
Single-cell transcriptomes of developing *Coccidioides*. **a**, *Coccidioides’* environmental life cycle. **b**, Each point represents a cell; positions represent transcriptomes projected according to UMAP, and colors report cluster membership. A subset of enriched Gene Ontology terms per cluster are shown; see S2 Table for the full list. **c**, Each row reports normalized expression from a cell-cluster-specific marker gene; color reports Z score. Column names report inferred cluster identities. **d**, Each column reports the distribution of the number of unique molecular identifiers detected in cells of the indicated cluster (grey box, interquartile range; black line, median; red diamond, mean). **e**, Data are as in **b** except that colors report *het-q1* expression relative to the maximum per cell.

## Results

We set out to use single-cell (sc) RNA-seq to identify candidate determinants of enteroarthroconidial development. As this technique has not previously been applied in filamentous fungi (although see related methods in [11–13]), we established a protoplasting protocol by which hyphal contents could be encapsulated for sequencing. We applied the approach to colonies of NR-166, an avirulent model of *C. posadasii,* cultured on solid medium until disjunctor cells had appeared but not yet emptied fully (six days after inoculation; see Fig 2a). From replicates of colony harvest, protoplasting, and sequencing, we confirmed reproducibility of expression levels by pseudobulk analysis (S1 Fig) and ultimately amassed profiles of 104,990 cells, with an average of 8,637 transcripts and 534 genes detected per cell. From the data, graph-based clustering in Monocle3 identified four transcriptomically distinct cell clusters (Fig 1b).

**Fig 2.**
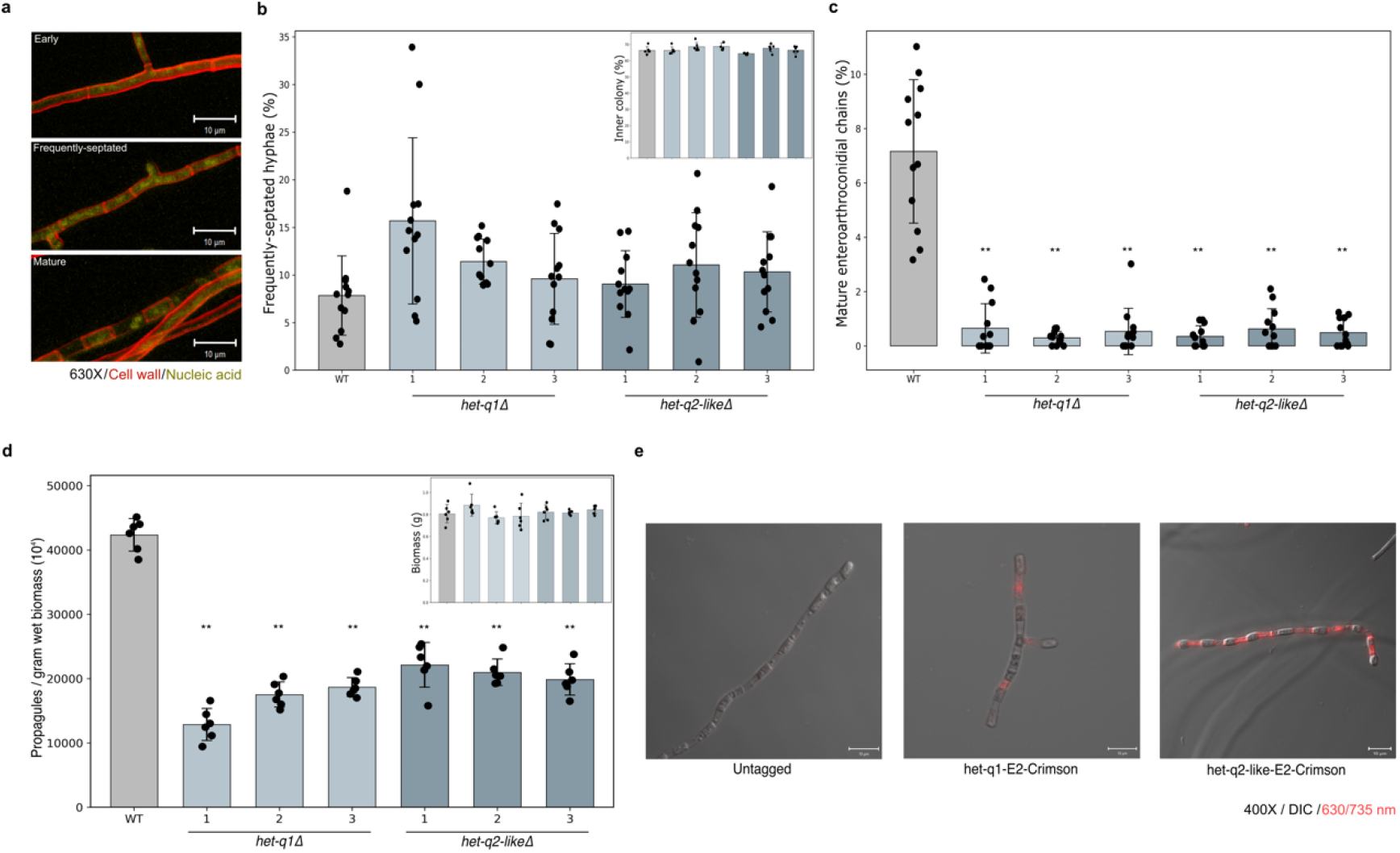
*het-q1* and *het-q2-like* drive enteroarthroconidial development. **a**, Representative images of wild-type (WT) *C. posadasii* aerial hyphae in the indicated stages of enteroarthroconidial development stained for cell wall (Calcofluor white, red) and nucleic acid (Thiazole orange, yellow). **b-c,** In the main plots, the *y*-axis reports the proportion of aerial hyphae falling into the indicated categories in WT and *het-q* mutants; points report individual replicates and bar height reports the average. **, Wilcoxon *p* < 0.01 in a comparison against WT. In the inset of **b**, column order is as in the main figure; the *y*-axis reports the percentage of a given colony visible as aerial hyphae. **d**, In the main plots, data are as in **b-c** except that the *y*-axis reports counts of disarticulated propagules normalized for colony biomass. In the inset, column order is as in the main figure; the y-axis reports colony biomass. **e**, Representative images of mature enteroarthroconidial chains in WT (Untagged) and isogenic strains expressing C-terminal E2-Crimson-tagged *het-q1* or *het-q2-like,* as a composite of images from transmitted light and the far-red fluorescent channel; see S2 Fig for all imaging data.

For insights into cluster identities, we inspected marker genes highly expressed in a given cluster relative to the others (S1 Table and Fig 1c). In one cluster we noted, among the markers, orthologs of regulators (*mst1*, *dbf2, sid1*) [3,14] and effectors (*aspA-E, bud3, bud4*, *rdi1, rho4*) [15–17] of septation, consistent with an identity as immature, septating aerial hyphae (Fig 1c, Aerial hyphae). Markers of a second cluster included orthologs of the cell death executioner genes *het-q1* and *het-s* [6,7], and we hypothesized that this cluster constituted disjunctor cells destined to die; it also expressed orthologs of ubiquitination and autophagy genes (*doa4, snf7*, *atg6-7*) [14,18,19] (Fig 1c, Disjunctor cells) and was distinguished by uniquely low RNA content (Fig 1d). The third cluster’s markers included orthologs of factors annotated in conidial maturation (*con6*) [20], cell wall synthesis (*hypA*, *kre9, wsc*) [21–23], and melanization (the tyrosinase D8B26_006956), consistent with an identity as enteroarthroconidia (Fig 1c, Enteroarthroconidia). A fourth cluster expressed orthologs of the vegetative growth repressor *flbA* [24] and conidiophore regulators (*medA*, *velB, mst1*) [3,25], suggestive of substrate-level mycelium initiating development (Fig 1c, Substrate mycelium). Unbiased functional analyses echoed these trends (Fig 1c and S2 Table); they also revealed enrichment of translation gene orthologs among markers of the putative enteroarthroconidial cluster (consonant with profiles of purified *C. posadasii* spores [26]; S3 Table), and transporter gene orthologs as markers of the putative substrate mycelium cluster, a signpost of nutrient depletion in aging colonies [27]. Together, these results underscore the power of scRNA-seq for elucidating candidate molecular signatures of *Coccidioides* cell identity.

As a validation test case for inferences from scRNA-seq, we considered the potential role of regulated cell death. We focused first on the *C. posadasii* ortholog of the *Podospora* gasdermin-like executioner *het-q1* [7], D8B26_007977, for which the highest-expressing cells fell in the putative disjunctor cell cluster, though more cells in other clusters expressed it at low levels (Fig 1e). We generated *het-q1* deletion mutants in the virulent wild-type Silveira *C. posadasii* background, cultured them on solid medium to the six-day timepoint, and investigated their morphology via quantitative microscopy, alongside wild-type controls (Fig 2a). In aerial hyphae sampled from wild-type, on average 80% of the biomass was in early stages of development; of the remainder, roughly half the hyphae were in the frequently-septated stage that precedes spore formation, and half were in a mature stage with thick-walled enteroarthroconidia and emptying disjunctor cells (Fig 2b-c). Relative to this profile, in *het-q1* mutants, the prevalence of mature chains in aerial hyphae dropped by four-to eight-fold (Fig 2c). No such effect of the *het-q1* mutation was detectable on colony growth or aerial hyphal formation or septation (Fig 2b and 2d), attesting to a specific function for *het-q1* in late enteroarthroconidial maturation. To assess the function of *het-q1* in dispersal, we cultured mutant colonies for eight days to advance development, and we then quantified their propagules (spores and incompletely fragmented spore chains) liberated by mechanical disruption, finding two-fold less disarticulation in the *het-q1* mutant relative to wild-type (Fig 2d).

Next, as *Podospora* Het-q1’s C-terminal inhibitory domain is cleaved by Het-q2, a serine protease encoded next to *het-q1* in the genome [7], we knocked out the syntenic homolog in *C. posadasii*, the annotated CHAT protease D8B26_007976, which we refer to as *het-q2*-*like*. The resulting mutants phenocopied their *het-q1* analogs in development and dispersal assays (Fig 2b-d). We conclude that the *het-q* genes are required for enteroarthroconidial chain maturation in *C. posadasii*. Imaging of wild-type *C. posadasii* expressing C-terminally-tagged *het* gene alleles confirmed protein localization in disjunctor cells, dovetailing with their predicted function in this cell type (Fig 2e and S2 Fig). In some chains the Het-q1 tag could also be detected at disjunctor cell borders and in enteroarthroconidia, suggestive of translocation between the segments (S2 Fig).

## Discussion

In this work, we have laid out a molecular model of cell types in *C. posadasii* and validated it with the discovery of *het-q* genes as determinants of enteroarthroconidial development. Advanced public health efforts to limit *Coccidioides’* transmission will rest on discoveries like ours of the mechanisms underlying spore production. The annotated protease that we characterize here, Het-q2-like, may be a particularly appealing potential target for small-molecule interventions to block *Coccidioides’* infectivity.

We propose that the *het-q* pathway, previously implicated in immune response and allorecognition in other fungi [28], mediates the regulated death of disjunctor cells in *Coccidioides* hyphae for a dual purpose: to recycle molecular components to fuel spore assembly [29] and to facilitate spore disarticulation. Such a model parallels recent discoveries implicating autophagy and ferroptosis in death and appressorium formation by multicellular *Magnaporthe* spores [9,10]. The emerging picture is one in which regulated cell death can play key developmental roles in fungi with complex life cycles, analogous to its well-studied biology in plants and animals. Our insights and approach lay the foundation for exploration of this principle across the fungal kingdom.

## Materials and methods

### Culturing and protoplasting for scRNA-seq

For a given replicate (n=3), 20 µL of a frozen stock of *Coccidioides posadasii* NR-166 enteroarthroconidia was inoculated onto 9 mL of 2X glucose-yeast-extract solid medium (2X GYE; 20 g/L glucose, 10 g/L yeast extract, 15 g/L BD Difco™ agar) in a 60 x 15 mm polystyrene petri dish and incubated at 30°C for one week. Following incubation, a 20 x 20 mm portion of mycelium biomass was retrieved using a cell scraper, placed into 3 mL of 0.01% Tween-80 in a 50 mL Falcon tube, and vortexed for 30 seconds. 20 µL of this biomass slurry was inoculated and evenly spread onto a plate of 2X GYE solid medium overlaid with cellophane (Neta Scientific Inc, RP1-1090) followed by incubation at 30°C for 6 days for subsequent protoplast generation. Following incubation, culture samples were first taken to examine using brightfield microscopy to check for developmental progress wherein hyphae were exhibiting frequent septation and had not reached a stage of mature enteroarthroconidial chain disarticulation. Next, using forceps, the biomass was peeled away from the cellophane and placed into 40 mL of protoplasting enzyme buffer (1.2 M Magnesium sulfate heptahydrate in 50 mM monobasic sodium phosphate buffer, pH = 5.0-5.5 adjusted with dibasic sodium phosphate heptahydrate, 50 mg/mL VinoTaste® Pro (Novozymes), centrifuged at 3000 *g* for 10 minutes, and filtered through a 0.45 µm PES filter (Celltreat, CT229709)). The biomass was shaken at 30°C and 80 rpm for three hours until protoplasts had formed (see S1 Movie). Protoplasts were filtered through two layers of 22-25 µm Miracloth (Sigma Aldrich, 475855) and collected into a 50 mL Falcon tube. Next, the protoplasts were slowly overlaid with 5 mL of 0.4M sorbitol-tris buffer (0.4M ST; 0.4M sorbitol, 50mM Tris, pH8)) and centrifuged in a swinging bucket rotor at 1200*g* with brakes at level 4 for 10 minutes. Protoplasts were then collected from the interphase, carefully mixed with 30 mL of 1M ST buffer (1M ST; 1M sorbitol, 50mM Tris, pH8.0), and incubated on ice for 15 minutes. Removal of digested cell wall fragments from protoplasts was done by centrifugation at 1200*g* at 4°C for ten minutes, followed by discarding of the supernatant, resuspension in 30 ml ST buffer, recentrifugation, and a final discard of the supernatant. The protoplasts were counted using a hemocytometer (Chemglass Life Sciences, CLS-4207), resuspended in 0.4M ST buffer to a final concentration of 5000 protoplasts/µL, and submitted for single-cell RNA-seq at the UC Berkeley Functional Genomics Lab, where protoplasts were encapsulated into gel bead-in-emulsions using the 10X Genomics Chromium Controller, followed by 3’ single-cell RNA-seq library preparation following the standard protocol (10X Genomics Chromium GEM-X Single Cell 3′ v4 Reagent Kit User Guide Chemistry). Libraries were sequenced using the NextSeq 2000 P2 platform (100SR/10X/400M reads). Replicates yielded transcriptomes for, respectively, 35,053 cells with a mean of 3,435 reads per cell; 42,423 cells with a mean of 11,786 reads per cell; 27,695 cells with a mean of 10,691 reads per cell.

### scRNA-seq read-mapping

A reference transcriptome was first generated using the 10X Genomics Cell Ranger mkref function with the *Coccidoides posadasii* strain Silveira genome and gene annotation from [30] (GenBank, BioProject PRJNA664774) as the inputs. Transcriptomes from a total of 105,171 cells were aligned to the reference transcriptome from the three biological replicates. For quality control, genes were retained for downstream analysis if they were detected in at least three cells; cells with fewer than 50 genes, fewer than 100 UMIs, or more than 2500 UMIs were eliminated from analysis, to account for cases of degraded cells and doublet/multiplet cells that were captured during GEM encapsulation, resulting in a final analyzable data set of 104,990 cells.

### scRNA-seq normalization and cell clustering

Raw RNA count matrices with associated cell-level metadata were combined from each replicate into a single cell_data_set within Monocle3 (v.1.4.26). Cell-specific size factors were utilized for library-size normalization for each dataset, followed by log-transformation and scaling during the initial preprocess_cds step implemented with Monocle3. Principal component analysis was performed using the first 20 principal components, followed by UMAP dimensionality reduction. Cells were then clustered using Leiden community detection by the cluster_cells() function default setting, using the cell-neighbor graph built from the UMAP reduction and a resolution of 2.35 × 10^-6^.

### scRNA-seq marker gene identification

Marker genes associated with each Monocle3-defined cell cluster were identified using the Seurat package’s FindAllMarkers function (Seurat v.5.4.0). For this purpose, RNA expression data was first normalized using the LogTransformation function, by which RNA counts for a given gene were divided by the total RNA counts for a given cell, followed by multiplication of a scale factor of 10,000 and subsequent log-transformation. Marker genes identified with positive average log_2_ fold-change values relative to the mean expression of all three other clusters and expressed in more than 1% of cells were retained. Significance testing was performed using the Wilcoxon rank-sum test and the resulting *p*-values were adjusted for multiple-testing using a Bonferroni correction. Marker genes with adjusted *p*-values < 0.05 were retained for analysis. Gene annotation was via MycoCosm and FungiDB.

### scRNA-seq UMAP overlay with *het-q1* expression

To visualize cells exhibiting the highest relative expression of *het-q1* in the scRNA-seq data set, cell-specific size factor-adjusted expression was overlaid onto the Monocle3 UMAP. The normalized expression values then were scaled to the maximum normalized expression value found across cells. Cells expressing less than 1% of the maximum scaled expression were plotted as grey points, and cells expressing *het-q1* up to the highest scaled maximum expression were highlighted by a color gradient and proportional point size. Points were plotted onto the UMAP graph in ascending order of scaled expression values for ease of viewing the highest-expressing cells.

### Functional-genomic analysis of marker genes from scRNA-seq clusters

Gene Ontology (GO) enrichment was performed independently on each Monocle3-derived cell cluster in turn as follows. Cluster marker genes identified by Seurat were used as the foreground genes. Gene identifiers were matched to GO annotations from the *Coccidoides posadasii* strain Silveira 2022 GAF file [30]. The background gene dataset consisted of all genes that were detected in at least 10 cells in the RNA count matrix. GO terms associated with fewer than five background genes were excluded from analysis. For each GO-term, a 2 × 2 contingency table was constructed comparing the number of foreground genes annotated with the term to the corresponding numbers of genes without the annotation in the foreground and background sets, with significance of enrichment assessed by one-sided Fisher’s exact tests. *p*-values were adjusted for multiple testing independently within each cluster using the Benjamini-Hochberg false discovery rate (FDR), and GO terms with an adjusted *p*-value < 0.05 were considered significantly enriched.

### Functional-genomic analysis of bulk RNA-seq

As a gold standard for comparison against scRNA-seq data, we evaluated expression differences between immature and fully differentiated *C. posadasii* cell types via bulk RNA-seq as follows. RNA-seq reads from two day-old hyphae and mature, purified enteroarthroconidia from *C. posadasii* Silveira[26] were retrieved from GEO (accession GSE279421). We used three replicates from each condition to generate a library size-normalized gene-by-sample count matrix. We then used DESeq2 (v.1.46.0) to assess expression differences between enteroarthroconidia and hyphae. *p*-values were adjusted for multiple testing using the Benjamini-Hochberg method, and genes were considered differentially expressed if they exhibited an adjusted *p*-value < 0.05 and abs(log_2_(fold-change)) > 0.5. The set of genes meeting these criteria that were more highly expressed in hyphae than in enteroarthroconidia and, separately, the analogous set more highly expressed in conidia, was then used as the foreground in GO enrichment analyses as above (see Functional-genomic analysis of marker genes from scRNA-seq); the background in each case comprised all genes input from the count matrices.

### Constructing gene knockouts

CRISPR/Cas9-mediated whole gene knockouts of *het-q1* and *het-q2-like* in the wildtype *C. posadasii* Silveira background were generated as described [31], using the Alternate Protocol 1 and 3 for gene deletion construct design and transformation of *Coccidioides* in a BSL-3 laboratory setting starting from frozen protoplasts, with minor adjustments as follows. For the DNA repair construct for *het-q1*, 203 bp upstream and 231 bp downstream sequences from the start and stop codon sites, respectively, were used as flanking sequences of the nourseothricin antibiotic resistance marker (NAT) cassette (from pDIV296, Addgene plasmid #204687, a gift from Uffe Mortensen). For *het-q2-like*, flanking sequences were 228bp upstream and 210bp downstream sequences from the start and stop codon sites, respectively. Constructs were synthesized by Twist Biosciences (South San Francisco, California) as gene fragments, amplified using Q5® High-Fidelity DNA Polymerase (NEB, Ipswich, MA, USA; #M0491S), cloned into the pCR™-Blunt II-TOPO® vector and then transformed into competent *Escherichia coli* from the Zero Blunt® TOPO® PCR Cloning Kit (Invitrogen/Thermo Fisher Scientific, Waltham, MA, USA; #K2800-J10). Plasmids were purified via QIAprep Spin Miniprep Kit (Qiagen #27106) from single *E. coli* colonies and sequenced by Plasmidsaurus (Louisville, KY) for confirmation of the construct sequences. Correct constructs were re-amplified from a given purified plasmid stock using Q5® polymerase to generate DNA repair templates, and the resulting PCR products were purified using the NucleoSpin Gel and PCR Clean-up kit (Macherey-Nagel, Düren, Germany; #740609.250). 2 µg of this DNA was used for each transformation together with the Cas9 ribonucleoprotein (RNP) complex. 5’ and 3’ single-guide RNAs (TCATTGAGAACCCAACACGA TGG) and (TCGACAGGATGCGTTTGGCG AGG), respectively were designed for the *het-q1* RNP complex. 5’ and 3’ single-guide RNAs (TCTCAAACCGGTCTCTGAGC CGG) and (CGTTGCCAGTGCCCTCCACC AGG), respectively, were designed for the *het-q2-like* RNP complex. Guide designs used CRISPOR. To confirm genomic integration in putative transformant colonies that survived three passages on selective media, diagnostic PCR checks for *het-q1* included primer pairs for amplification of the region upstream of the 5’ homology arm used in the DNA repair construct (CATCAGGCAAAGTGTGATGGG) into the NAT cassette (ccaatccacaaagtgtgaaccc). PCR primers were used to amplify the region downstream of the 3’ homology arm used in the DNA repair construct (ATCCTGAAGTCATCTATGGTGACC) and into the NAT cassette (atgctgaagctatcgaagcc). Finally, we used primer pairs (CTAGAAGACGTCCAGTTGGC) and (CTGGTCTTCAGATTCTTCACCC) homologous to regions inside the native gene to confirm the absence of native *het-q1* in the transformants. We applied the same strategy for *het-q2-like* transformants. Primer pairs were used for amplification upstream of the 5’ homology arm used in the DNA repair construct (GGCAATTATGATCACCTGAATTGGG) into the NAT cassette (ccaatccacaaagtgtgaaccc). Amplification of the region downstream from the 3’ homology arm to inside the NAT cassette used primer sets (CAGAGTTGAACCCAGTGACC) and (atgctgaagctatcgaagcc), respectively. We used primer pairs (CCATATACCCGCTTGTAACCC) and (GAAAAGCGACTTGAGAGAACCG), to confirm the deletion of *het-q2-like* in the transformants. All primers were from IDT (Coralville, Iowa).

### Constructing fluorescently-tagged alleles

CRISPR/Cas9-mediated E2-Crimson fluorescent-tagging of *het-q1* and *het-q2-like* in the wildtype *C. posadasii* Silveira background were generated using protocols analogous to those for whole-gene knockouts (see Constructing gene knockouts); in each repair template, the location corresponding to the PAM site in the coding sequence was substituted with a silent mutation that eliminated the PAM site but maximized the similarity to the native codon with respect to codon usage. The *het-q1* construct consisted of the sequence starting from the 3’ sgRNA cut site used in our whole-gene knockout DNA repair fragments (185 bp upstream from the stop codon), followed by a hemagglutinin (HA) tag and a GA5 linker hinge region as described [32], tatccatatgatgtcccagattatgcc and GGAGCTGGTGCAGGCGCTGGAGCCGGTGCT, respectively. Next, the E2-Crimson fluorescent protein sequence was fused to the GA5 linker, followed by the short *trpC* terminator from *Aspergillus nidulans* and NAT cassette (see Constructing gene knockouts) and 231 bp of sequence downstream of the *het-q1* stop codon. The *het-q2-like* construct consisted of the sequence starting from the 3’ sgRNA cut site used in our whole-gene knockout DNA repair fragments (94 bp upstream from stop codon), followed by the HA-GA5-E2-Crimson-NAT cassette as above. Both constructs were synthesized by Twist Biosciences, cloned into TOPO vectors and into *E. coli*, then purified, sequenced, and re-amplified as above (see Constructing gene knockouts). 1 µg of DNA was used together with Cas9 RNP complexes for transformations. To confirm genomic integration of the E2-Crimson fluorescent protein in putative transformant colonies that survived passages on selective media, diagnostic PCR checks for *het-q1*-E2-Crimson-NAT included primer pairs for amplification from a region upstream of the 3’ sgRNA cut site (ctggcaggtacgaacaagg) through the HA-GA5-E2-Crimson cassette and end at the *trpC* promoter within the NAT cassette (ggcattgatgtgttgacctcc). Additionally, we confirmed integration of the NAT cassette through primer pairs starting from inside NAT (atgctgaagctatcgaagcc) and ending 294 bp downstream from the *het-q1* stop codon (ATCCTGAAGTCATCTATGGTGACC). Similarly for *het-q2-like-*E2-Crimson-NAT transformants, primer pairs starting upstream of the 3’ sgRNA cut site (atcagctatgagcatgcacg) through the HA-GA5-E2-Crimson cassette and end at the *trpC* promoter (ggcattgatgtgttgacctcc) were used for amplification. Additionally, we confirmed integration of the NAT cassette through primer pairs starting from inside NAT (atgctgaagctatcgaagcc) and ending 262 bp downstream from the *het-q2-like* stop codon (CAGAGTTGAACCCAGTGACC). All primers were from IDT (Coralville, Iowa).

### Common source stocks for phenotyping

Dispersal, development, growth, and imaging assays, detailed below, each used a design in which a culture of a given *C. posadasii* strain served as a common source of inoculum for all replicates’ worth of experimental cultures for that strain. To make these common source stocks, we prepared 3-4 60 x 15 mm polystyrene petri dishes, each containing 9 mL of 2X GYE solid medium; we added, as a point inoculum, a small volume of mycelium of a given strain (wildtype *C. posadasii* Silveira control or each isogenic *het-q1* or *het-q2-like* mutant in turn) from a previously propagated culture, using a pipet tip. The resulting stocks were retained for 2-6 months at 30°C and used as source material as described below.

### Biomass and propagule count assay

To quantify propagule counts from developing colonies of wildtype *C. posadasii* Silveira control or an isogenic strain bearing a whole-gene knockout mutation of *het-q1* or *het-q2-like*, for a given biological replicate we prepared each of three 60 x 15 mm polystyrene petri dishes with 9 mL of 2X GYE solid medium overlaid with cellophane. We retrieved a 20 x 20 mm portion of mycelium biomass from a stock of the respective strain using a cell scraper (see Common source stocks, above), placed it into 3 mL of 0.01% Tween-80 in a 50 mL Falcon tube, and vortexed for 30 seconds. Then 20 µL of the resulting slurry was evenly spread onto each of the three prepared cellophane plates. Plates were incubated at 30°C for eight days. Biomass from each sample was then harvested into a pre-weighed 50 mL Falcon tube containing 9 mL of 0.01% Tween-80. Tubes were subsequently weighed to determine the wet biomass of each sample after subtracting the weight of the tube and Tween-80 solution. Each sample was then shaken vigorously three times to disrupt the mycelial mass, followed by one minute of vortexing, and filtered through Miracloth into a new 50 mL Falcon tube. For quantification of disarticulated propagules, 250 µL of filtrate was mixed with 250 µL of 10% formalin and then counted using a hemocytometer. The resulting concentration was used to determine the total concentration based on the total filtrate volume. Final propagule counts were normalized to the corresponding wet biomass and reported as propagules per gram of wet biomass. The complete biological replicate procedure, from slurry formation through eight days of development to harvest and quantification, was carried out twice for each mutant strain, with an accompanying wild-type Silveira control in each case.

### Nucleic acid and cell wall stain microscopy

To visualize nucleic acid and cell wall in septa and enteroarthroconidial chains (Figure 2A), we inoculated and incubated an experimental plate of wildtype *C. posadasii* Silveira from a common source stock as above (see Propagule count assay), except that incubation of experimental plates was for six days. A sample of vertically-growing hyphal tissue was collected from the plate ∼5 mm from the edge using forceps. We placed it into a 2 mL screw-top tube containing a working solution of 2 µL of 10X Thiazole Orange nucleic acid fluorescent stain from the Yeast Live-or-Dye™ Fixable Live/Dead Staining Kit (Biotium, #31064) in 200 µL of 1X PBS. The sample was left to incubate for 30 minutes with light mixing every 10 minutes, followed by addition of 200 µL 10% formalin for tissue fixation. Next, 10 µL of Calcofluor White (M2R, 1 g/L, Evans blue, 0.5 g/L) cell wall stain (Millipore Sigma, 18909) was added to the sample and incubated for an additional 15 minutes. Next, we collected the tissue sample from the staining solution and mounted it onto a glass microscope slide in 10 µL of 1X PBS, overlaid with a cover slip, and sealed with clear nail polish for long-term storage. Imaging was performed using a Zeiss LSM-710 confocal microscope system and Zeiss ZEN Black imaging software at the UC Berkeley Biological Imaging Facility. The Z-stack and subsequent deconvolution imaging features were used to capture all focal planes in z-steps of 0.63 microns at 630X magnification. Calcofluor White fluorescence utilized 405/430 nm excitation/emission and Thiazole Orange utilized 488/533 nm excitation/emission, respectively. Finally, composite images were generated using the Zeiss ZEN Black software, with cell wall fluorescence reported as red and nucleic acid fluorescence reported as yellow.

### Quantitative microscopy of development

Septum and enteroarthroconidial chain development in aerial hyphae (Figure 2B-C) were quantified as follows. For a given biological replicate of wildtype *C. posadasii* Silveira control or a *het-q1* or *het-q2-like* mutant strain, we inoculated and incubated two technical replicate experimental plates from common source stocks as above (see Propagule count assay), except that incubation of experimental plates was for six days. A sample of vertically-growing hyphal tissue was collected from each plate ∼5 mm from the edge using forceps. Each was stored in a tube containing 250 µL of 10% formalin for tissue fixation. For microscopy, each fixed tissue sample was transferred into a new tube containing 200 µL of 1X PBS, followed by a fifteen-minute incubation after the addition of 10 µL of Calcofluor White cell wall stain as above (see Nucleic acid stain microscopy). We transferred each sample to a microscopy slide and captured images from three fields at 200X magnification as above (see Nucleic acid stain microscopy) using 405/430 nm excitation/emission to capture cell wall and septum-stained fluorescence. The complete biological replicate procedure was carried out twice for each mutant strain, with an accompanying wild-type Silveira control in each case.

Image analysis was as follows. We categorized the developmental state of each hypha in an image as infrequently septated, frequently septated, or a mature enteroarthroconidial chain. Utilizing ImageJ (v1.54), the Set Scale function was set to microns using a previously-overlaid scale bar from Zeiss ZEN Black software, and each image was traced along the total length of hyphae in each category in turn with the Segmented Line tool, using the ROI Manager Add [t] function to store measurements. After all hyphae were measured in a given category for the image, Image > Overlay > Flatten created a new image showing all hyphae with overlaid outlines in yellow and numbered for reference. Next, within the ROI Manager, the Measure function generated a file containing all measured hyphae in microns, which was saved as a .CSV table. Finally, for a given image, we tabulated the total microns of hyphae in each category in turn, and we normalized each such value by the sum across all three categories, expressed as percentages. After collation of the resulting values across all images for a given strain (from all microscopy fields, culture plates, and biological replicates), significance testing was performed using the Wilcoxon two-sided rank-sum test comparing the wild-type with each *het-q1* or *het-q2-like* strain in turn; the resulting *p*-values were corrected for multiple testing using the Benjamini-Hochberg method.

### Radial growth assay

For growth assays, for a given biological replicate of wildtype *C. posadasii* Silveira control or a *het-q1* and *het-q2-like* strain, for each of three technical replicates, we took a 5 mm circular plug of agar containing fungal culture from a common stock plate of the respective strain (see Common source stocks, above) and transferred it onto the center of a 100 x 15 mm petri plate containing 25 mL of 2X GYE solid medium. These experimental plates were incubated at 30°C and radial growth measurements were taken on day six using a ruler. The total radial growth diameter of a colony was measured in centimeters from one side of the leading growth edge to the opposing edge and logged. To quantify the proportion of the colony’s radial growth attributable to aerial hyphae, the inner diameter of mycelial growth was measured that was more visibly dense with vertical hyphal growth present. To calculate the percentage of radial growth that had developed into aerial hyphae, measurements were first converted from centimeters into millimeters, followed by subtraction of 5 mm from each measurement to exclude the length of the agar plug. Next, the diameter of the dense inner radial colony measurement was divided by the total radial growth measurement, and this ratio was multiplied by 100. The complete biological replicate procedure was carried out twice for each mutant strain, with an accompanying wild-type Silveira control in each case. Significance testing was as above (see Propagule count assay).

### Microscopy of tagged proteins

Visualization of E2-crimson-tagged Het-q1 and Het-q2-like protein localization in mature enteroarthroconidial chains was performed as follows. For a given biological replicate, wildtype *C. posadasii* Silveira control and *het-q1-*E2-crimson and *het-q2-like-*E2-crimson tagged strains were inoculated from common source stocks as above (see Propagule count assay) and the experimental plates were incubated for six days. A sample of vertically-growing hyphal tissue was collected from each plate ∼5 mm from the edge using forceps. Each was stored in a tube containing 250 µL of 10% formalin for tissue fixation. For microscopy, each fixed tissue sample was transferred into a new tube containing 200 µL of 1X PBS for rinsing. We transferred each sample to a microscopy slide and captured images from five fields at 200X magnification as above (see Nucleic acid stain microscopy) using transmitted light T-PMT and 630/735 nm excitation/emission to capture far-red fluorescence of the E2-crimson tagged proteins. The complete biological replicate procedure was carried out twice for each strain.

## Acknowledgements

The authors thank Jules Cho and Denise Schichnes for microscopy assistance; Bridget Barker, Lisa Couper, and Anita Sil for *Coccidioides* expertise; and Russell Vance, Louise Glass and members of the Brem lab for helpful discussions.

## Data availability

scRNA-seq data will be available in the Gene Expression Omnibus upon publication. Images, image quantification data, propagule counts, and colony diameter measurements are available on Figshare: 10.6084/m9.figshare.34265337.

## Supporting information

**S1 Fig**. **scRNA-seq cluster markers.** Each row reports results for one gene called as a cluster marker in *C. posadasii* scRNA-seq, exhibiting higher expression in the cells of a focal cluster than in the union of the other three. The gene and its annotation are reported in the first two columns, and the focal cluster is listed in the third column. p_val and p_val_adj report the nominal and Benjamini-Hochberg corrected significance, respectively, of the expression difference between the focal cluster and the other three. avg_log2FC reports the log2 of the ratio between the expression in the focal cluster and the mean expression across the cells of all other clusters. pct.1 and pct.2 report the percentage of cells in which the gene is detected in the focal cluster and in the union of all other clusters, respectively. Clusters 1, 2, 3 and 4 correspond to those labeled in Figure 1 of the main text as Substrate mycelium, Disjunctor cells, Aerial hyphae, and Enteroarthroconidia, respectively.

**S2 Fig**. **Het-q protein localization during development.** Each page reports results from imaging of enteroarthroconidial chains in one field of aerial hyphae from wild-type Silveira *C. posadasii* (WT) or an isogenic strain expressing an E2-Crimson-tagged allele of *het-q1* or *het-q2-like,* after six days of growth on solid medium, for the indicated biological replicate. On a given page, boxes show images captured from transmitted light (top left) and the far-red fluorescent channel (top right), and their composite (bottom left).

**S1 Table. scRNA-seq cluster markers.** Each row reports results for one gene called as a cluster marker in *C. posadasii* scRNA-seq, exhibiting higher expression in the cells of a focal cluster than in the union of the other three. The gene and its annotation are reported in the first two columns, and the focal cluster is listed in the third column. p_val and p_val_adj report the nominal and Benjamini-Hochberg corrected significance, respectively, of the expression difference between the focal cluster and the other three. avg_log2FC reports the log2 of the ratio between the expression in the focal cluster and the mean expression across the cells of all other clusters. pct.1 and pct.2 report the percentage of cells in which the gene is detected in the focal cluster and in the union of all other clusters, respectively. Clusters 1, 2, 3 and 4 correspond to those labeled in Figure 1 of the main text as Substrate mycelium, Disjunctor cells, Aerial hyphae, and Enteroarthroconidia, respectively.

**S2 Table. Functional genomics of cell-type-specific expression programs.** Each tab reports results of tests for enrichment of Gene Ontology (GO) terms among marker genes of a focal cell cluster from *C. posadasii* scRNA-seq, those inferred to represent aerial hyphae, disjunctor cells, enteroarthroconidia, and substrate mycelium, respectively. In a given tab, each column reports results for one term. The first two columns report the term and annotation. Marker_Genes_in_GO reports the number of genes called as markers of the focal cluster assigned to the term. Total_Genes_in_GO reports the total number of genes in the genome assigned to the term. OddsRatio reports the proportion of marker genes assigned to the term, relative to the analogous quantity for the genomic background. P_Value and Adjusted_P report nominal and Benjamini-Hochberg corrected significance, respectively, in a Fisher’s Exact Test for enrichment of term membership among the marker genes.

**S3 Table. Functional genomics of hyphae and enteroarthroconidia bulk expression programs.** Data are as in S2 Table, except that each tab reports results of tests for enrichment of GO terms among genes differentially expressed in a comparison between two bulk culture regimes of *C. posadasii:* hyphae two days after germination and purified enteroarthroconidia. Results from the sets of genes more highly expressed in enteroarthroconidia and hyphae are reported in tabs with the respective labels.

**S1 Movie. Protoplasting developing hyphae**. Shown are the results of imaging of a mycelial sample from a *C. posadasii* NR-166 colony grown on solid medium for 6 days, sealed onto a microscope slide with Vinotaste enzyme cocktail and allowed to protoplast. Micrographs were captured at 400X magnification in 15-minute intervals for a duration of 15 hours.

