## Supplementary material for "Genetics and genomics of enteroarthroconidial development in *Coccidioides*": Fig S2

### WT, Field 1, Replicate 1

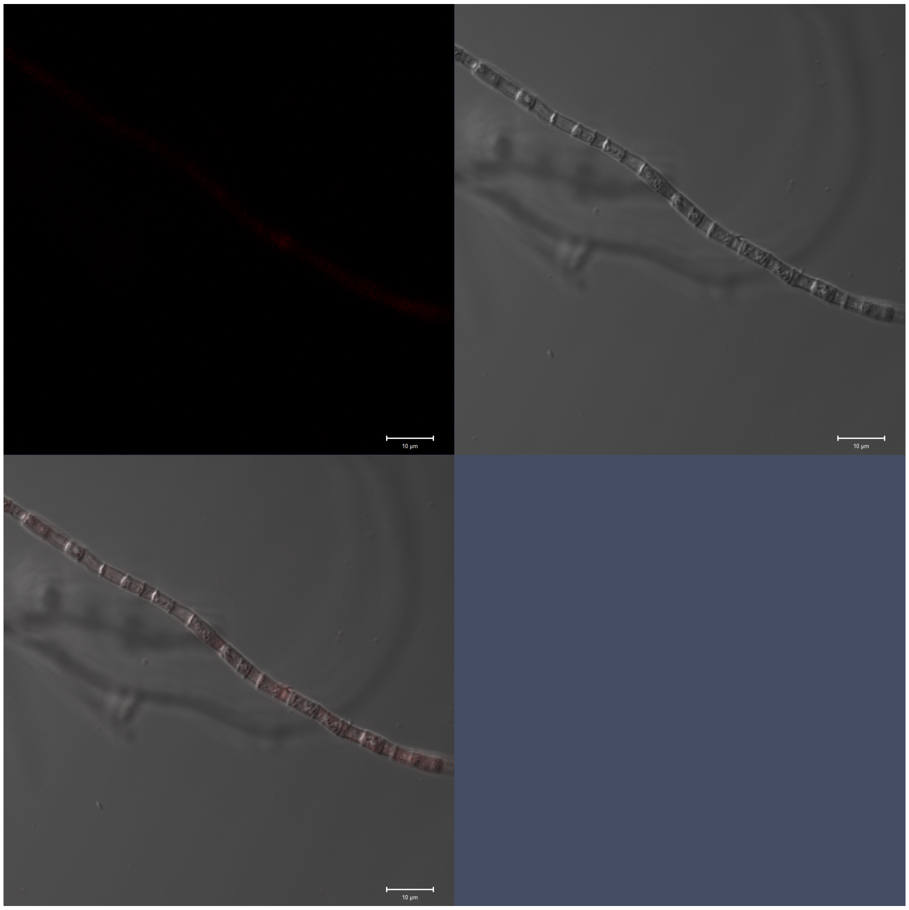

### WT, Field 2, Replicate 1

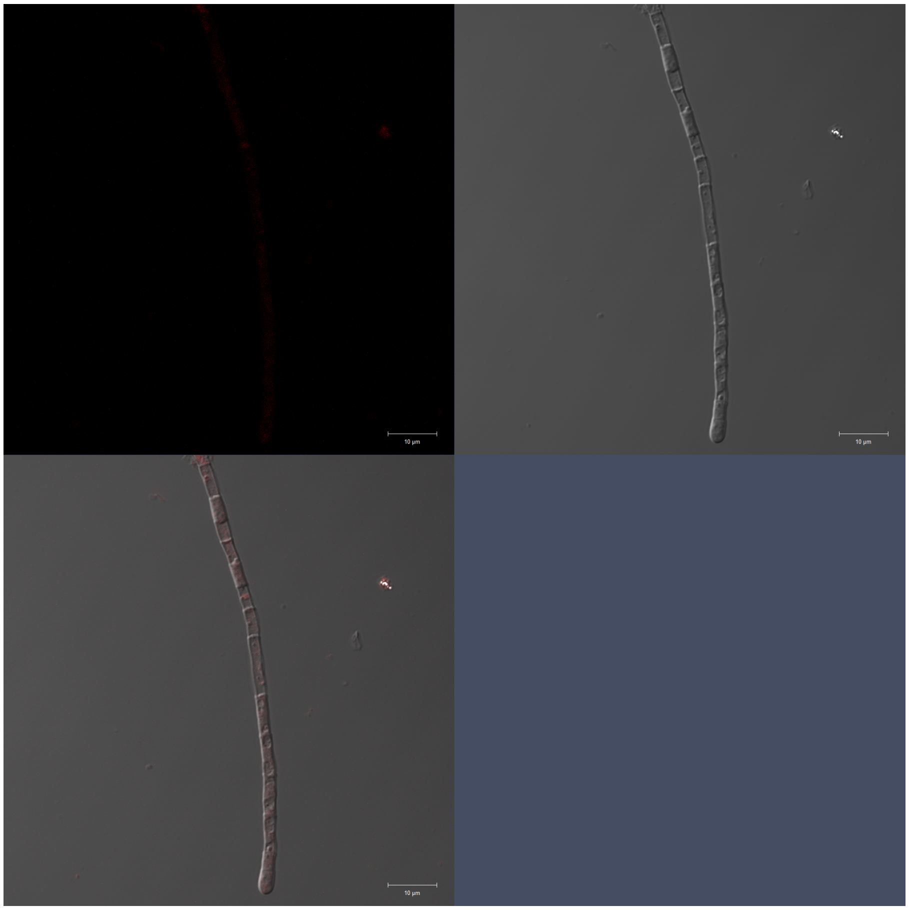

WT, Field 3, Replicate 1

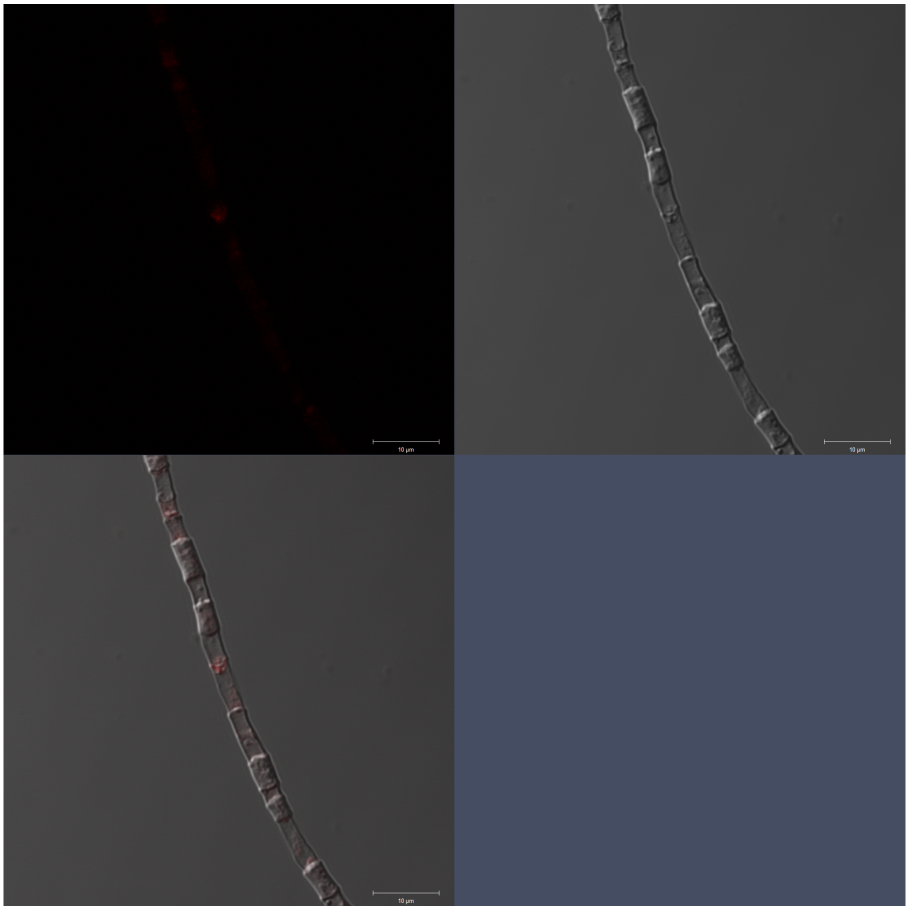

WT, Field 4, Replicate 1

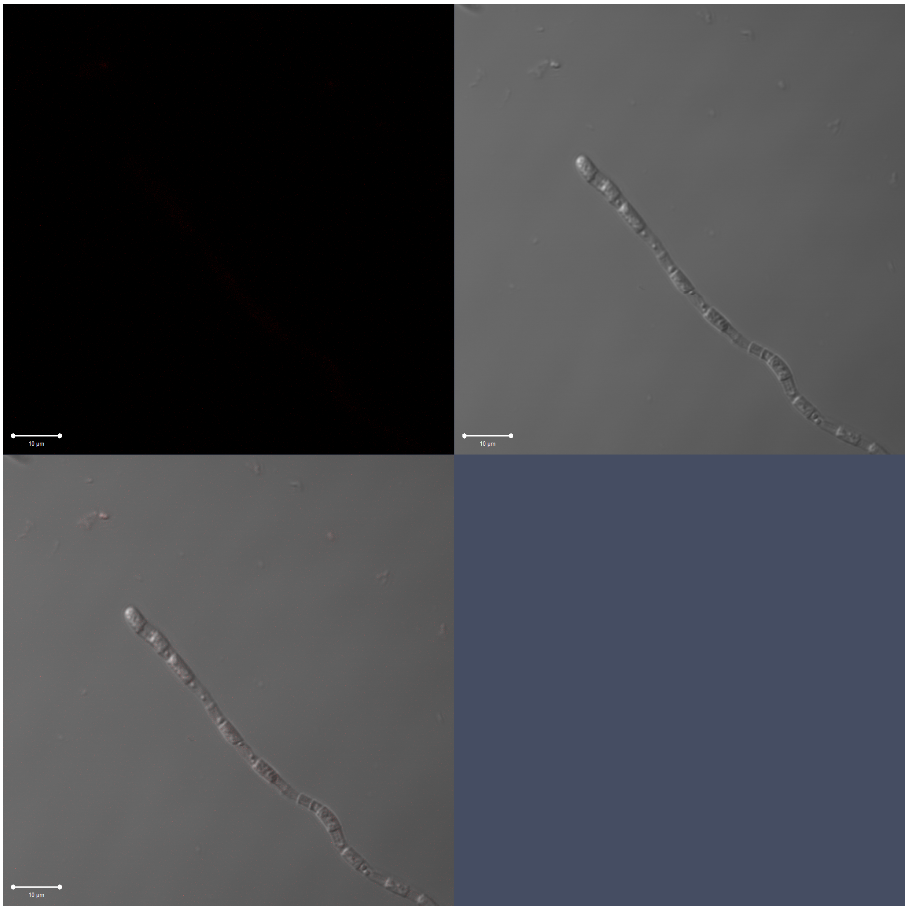

WT, Field 5, Replicate 1

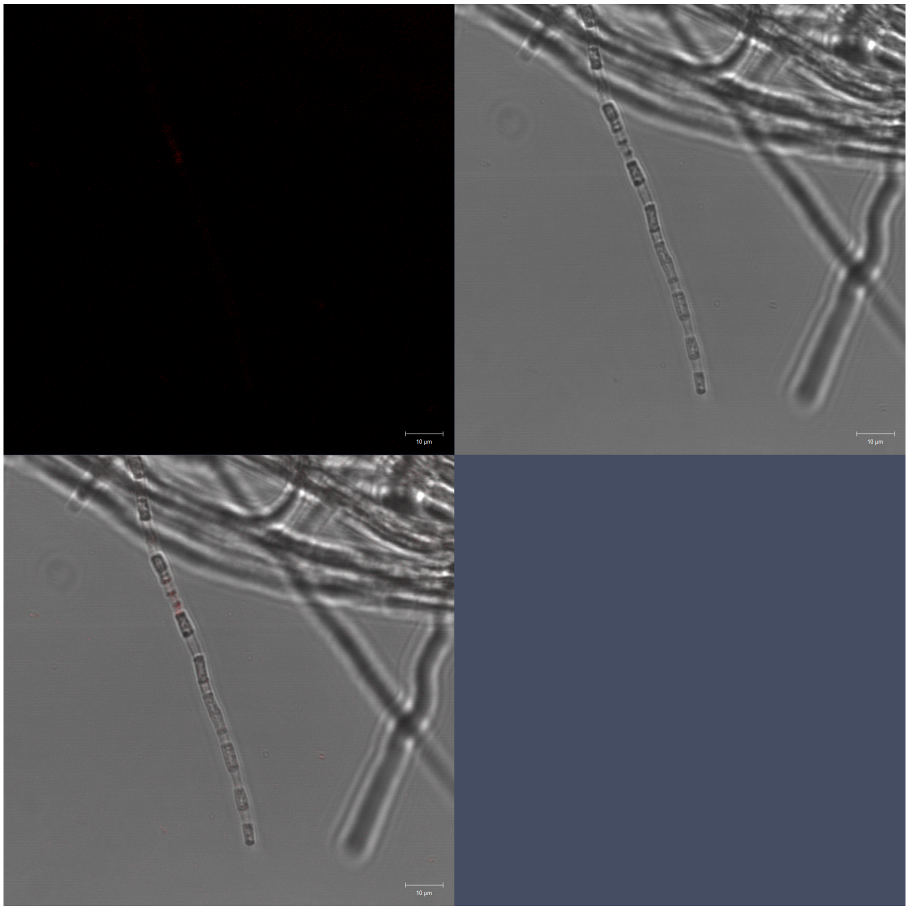

WT, Field 1, Replicate 2

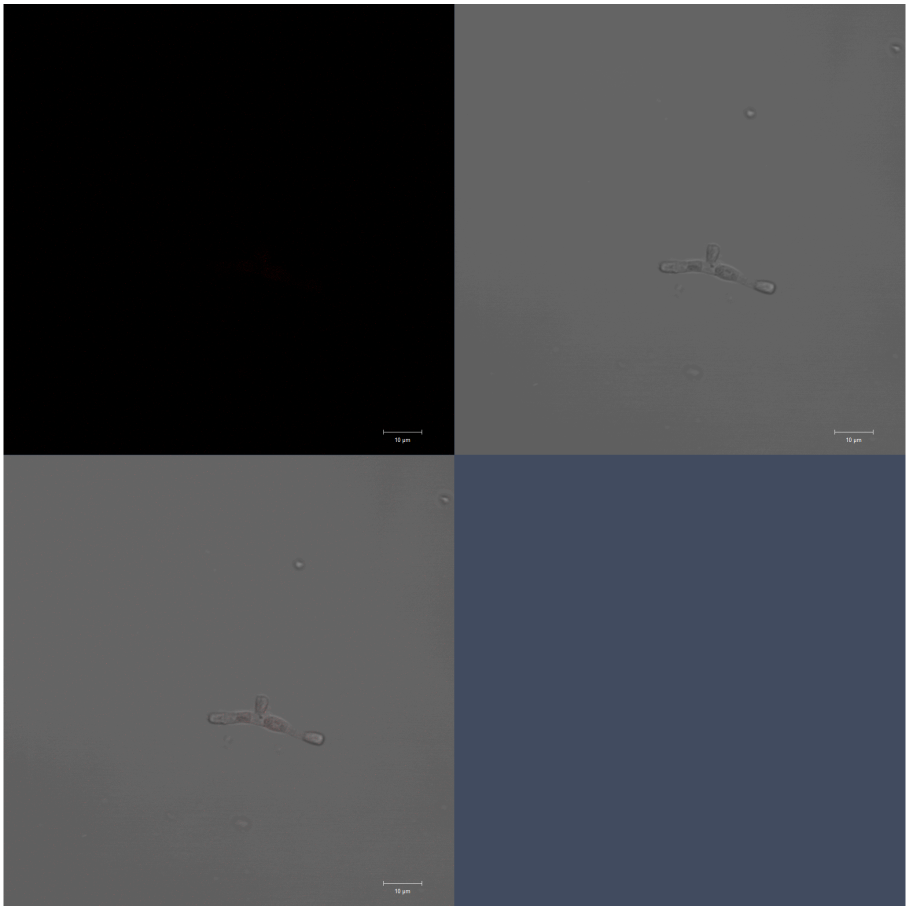

WT, Field 2, Replicate 2

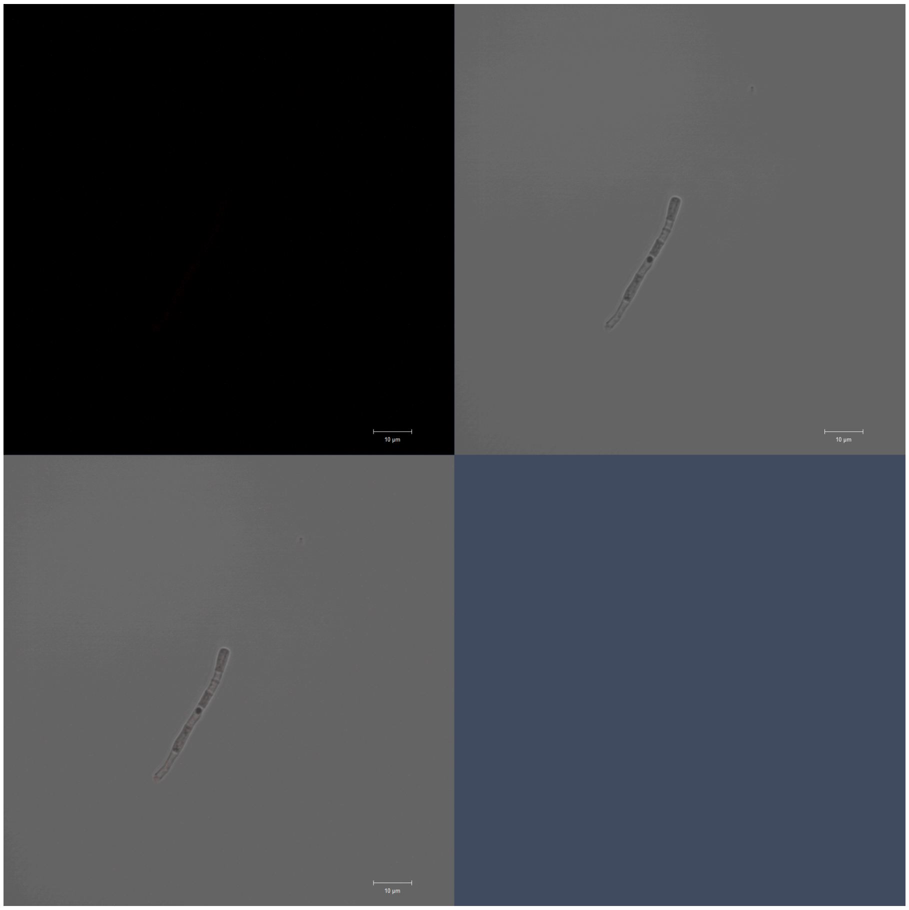

WT, Field 3, Replicate 2

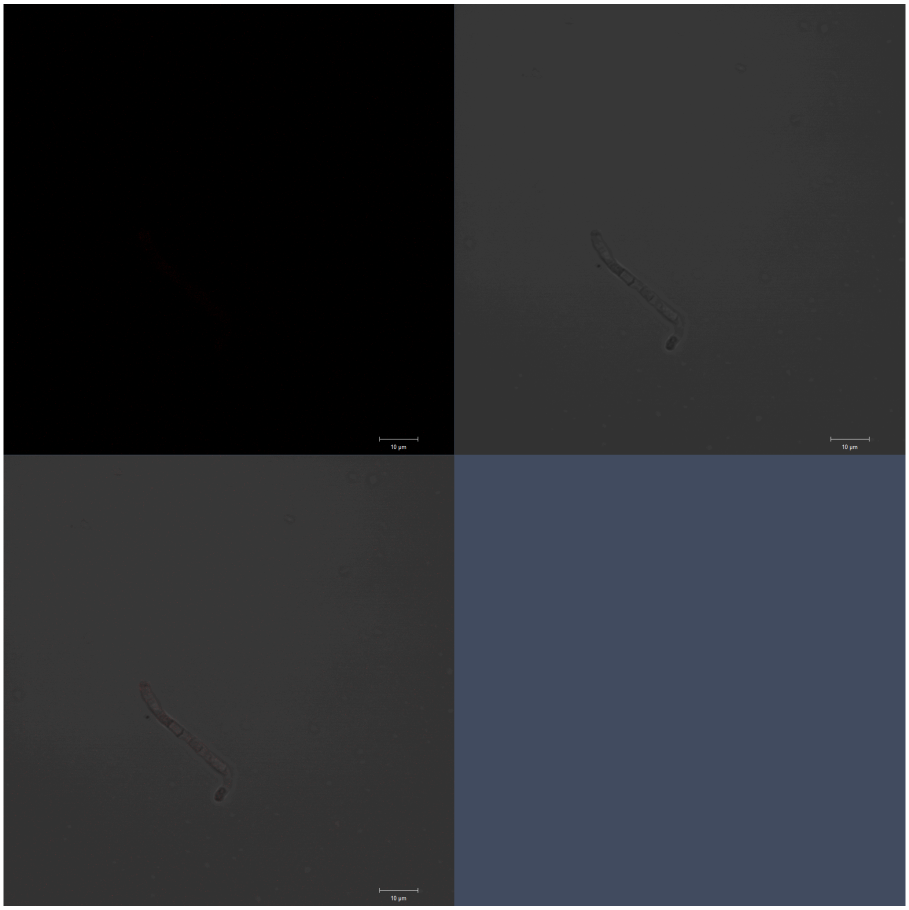

WT, Field 4, Replicate 2

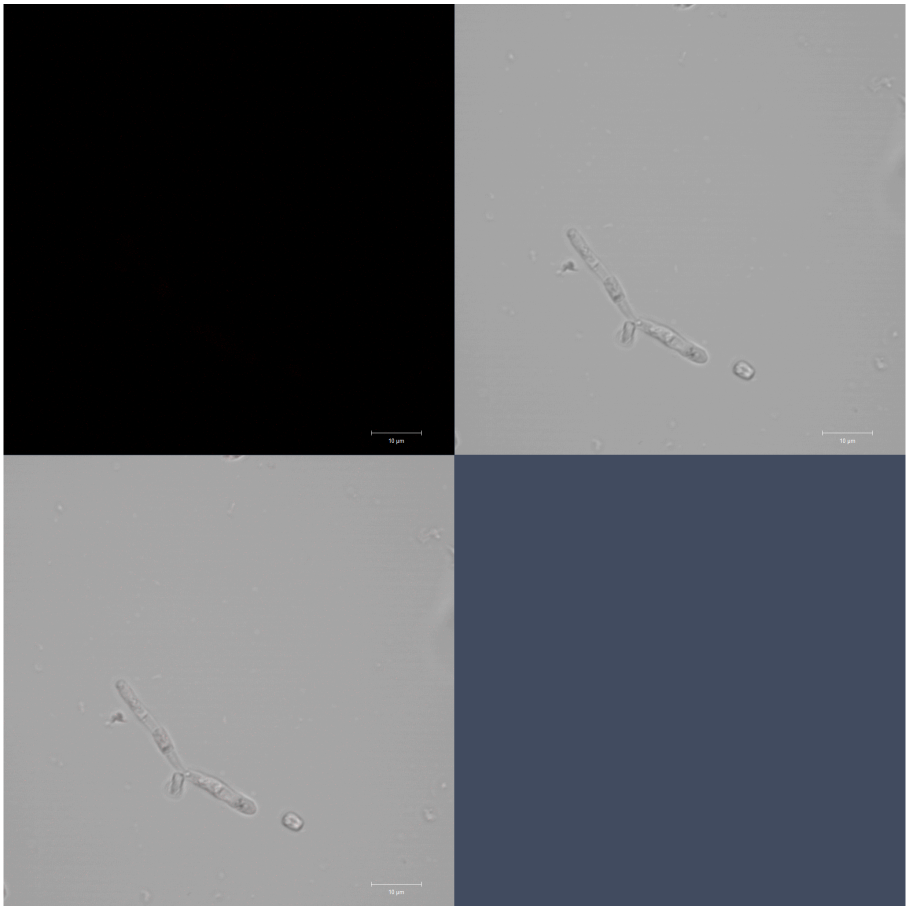

WT, Field 5, Replicate 2

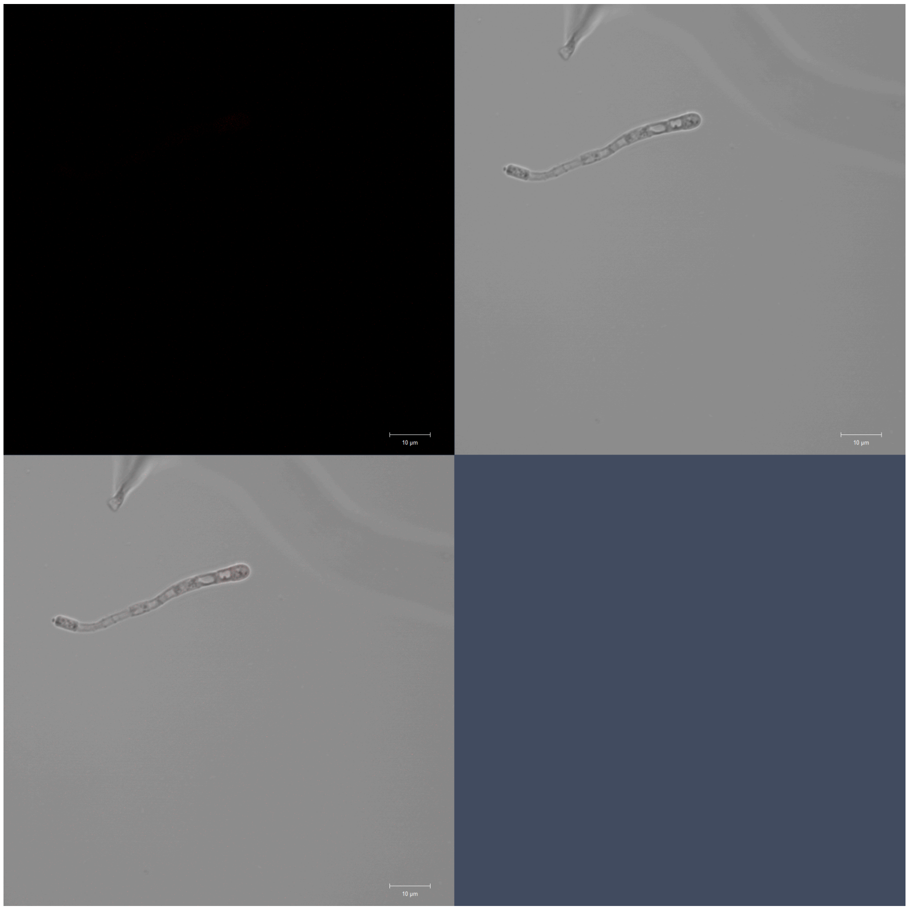

het-q1, Field 1, Replicate 1

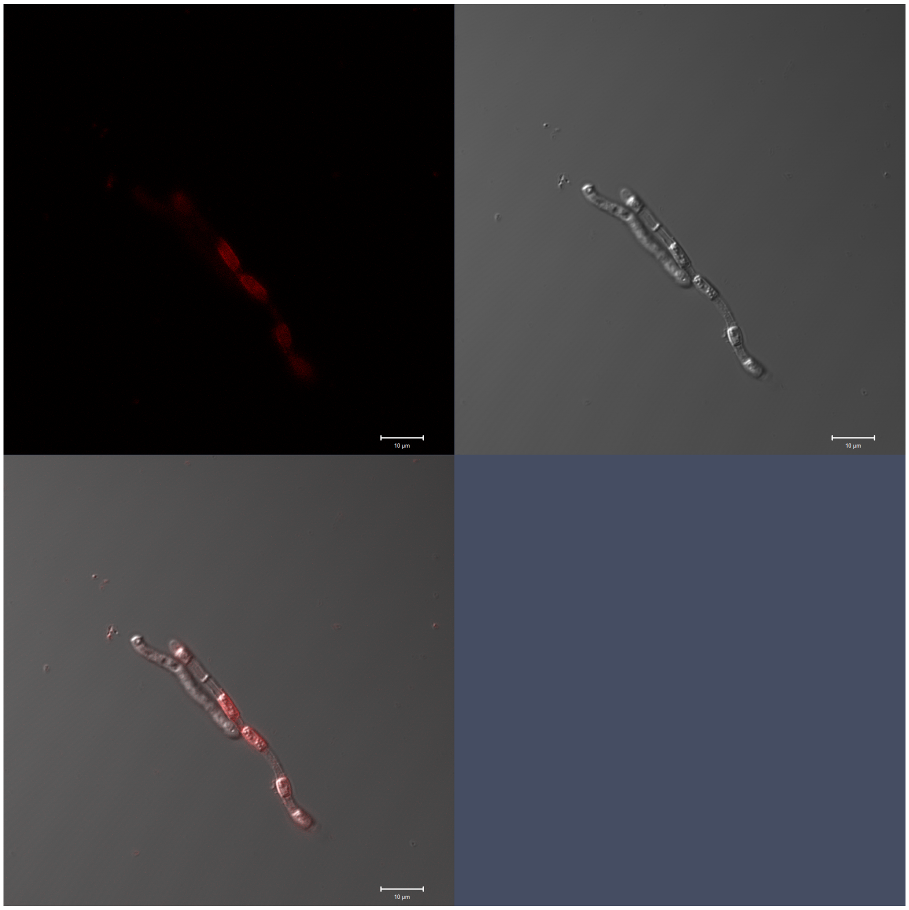

het-q1, Field 2, Replicate 1

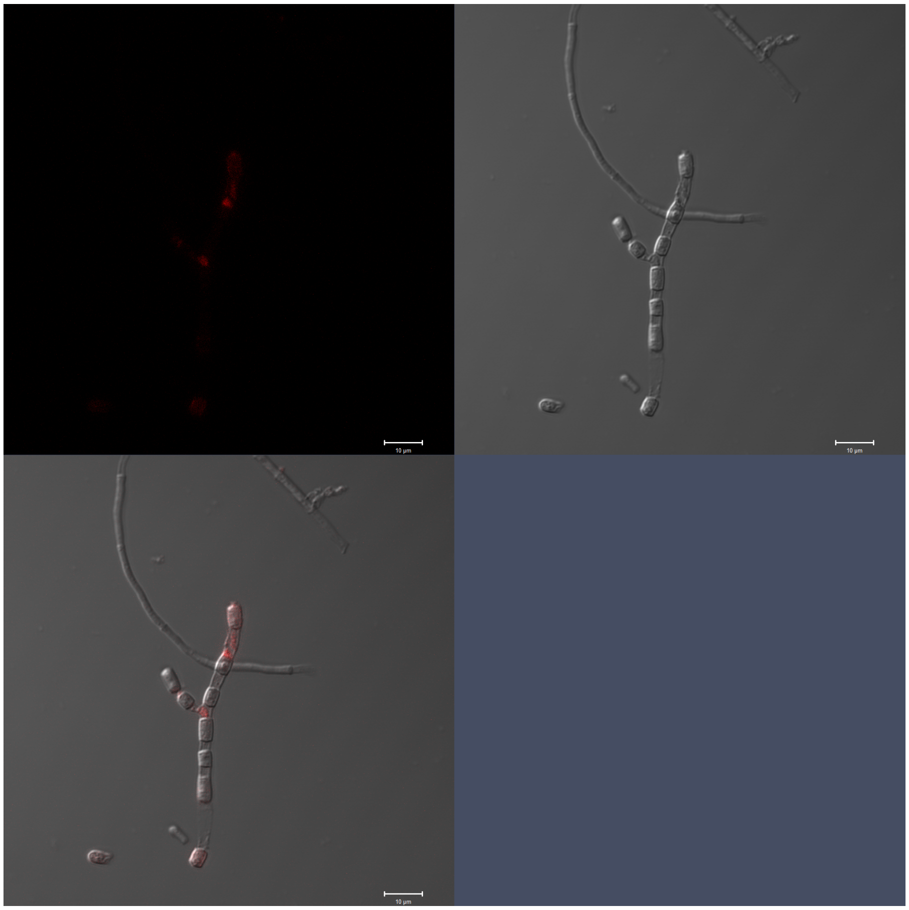

het-q1, Field 3, Replicate 1

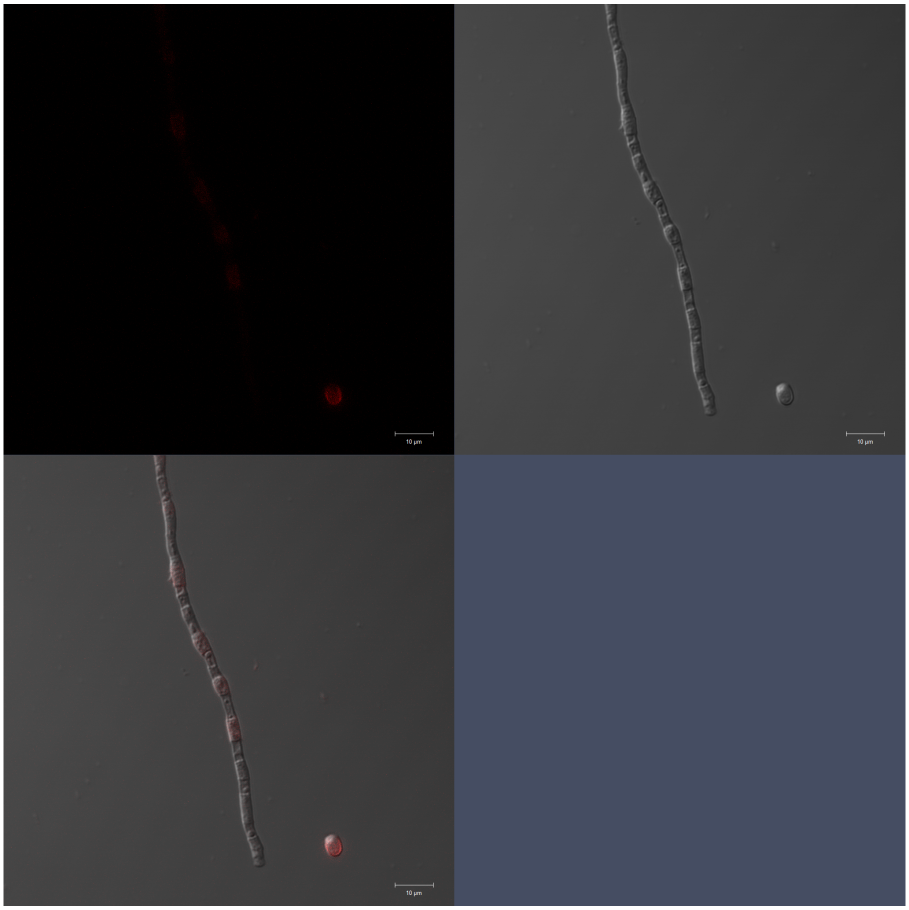

het-q1, Field 4, Replicate 1

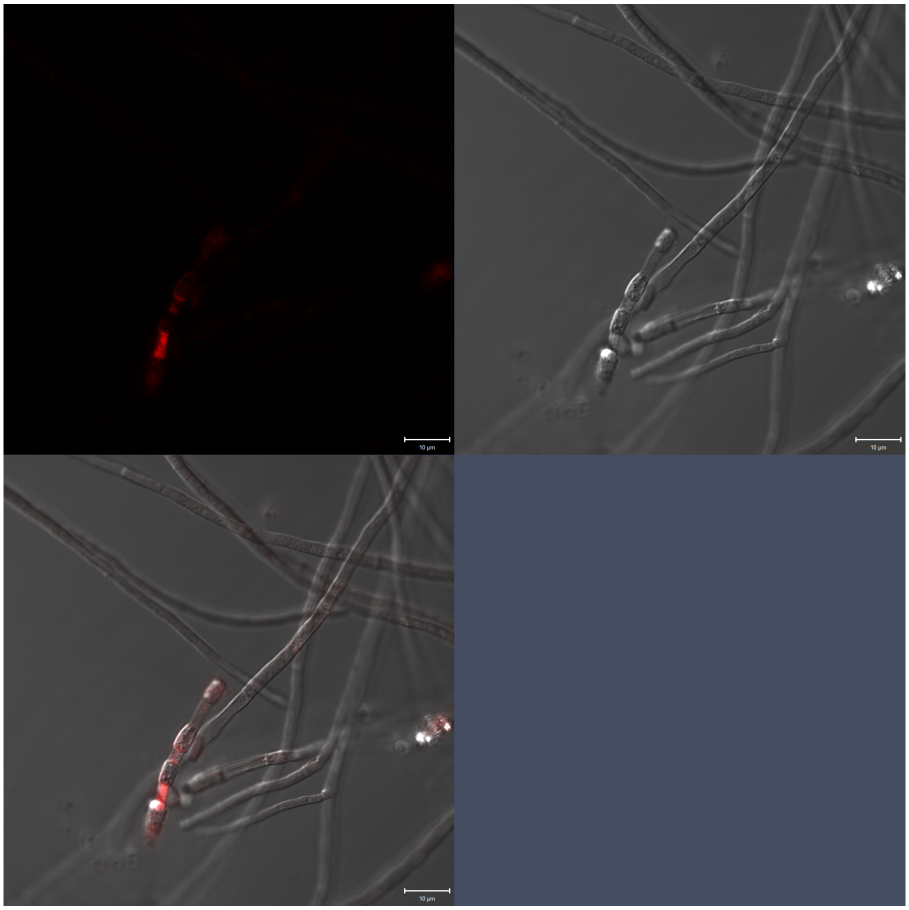

het-q1, Field 5, Replicate 1

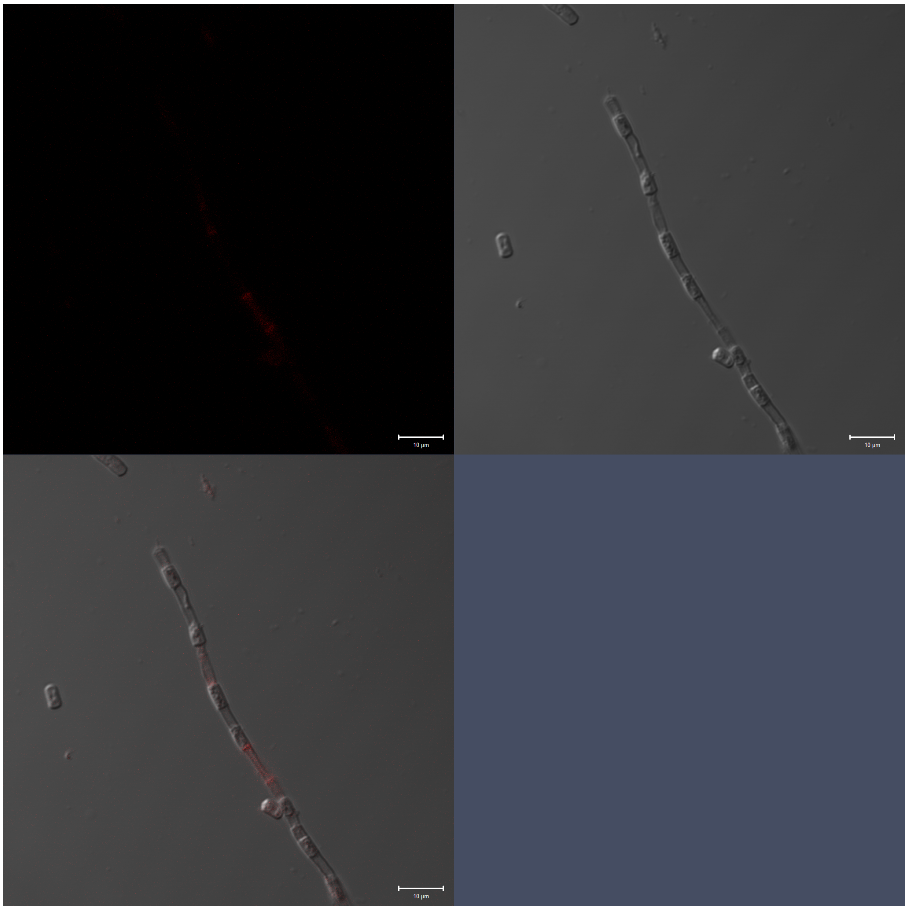

het-q1, Field 1, Replicate 2

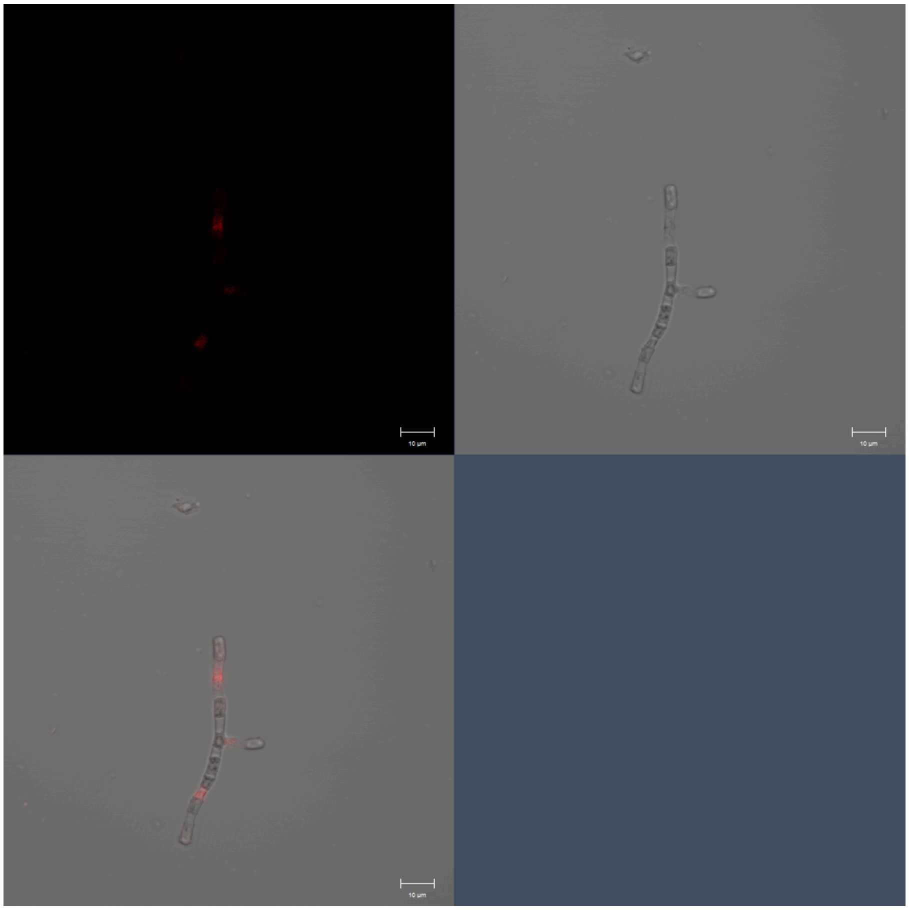

het-q1, Field 2, Replicate 2

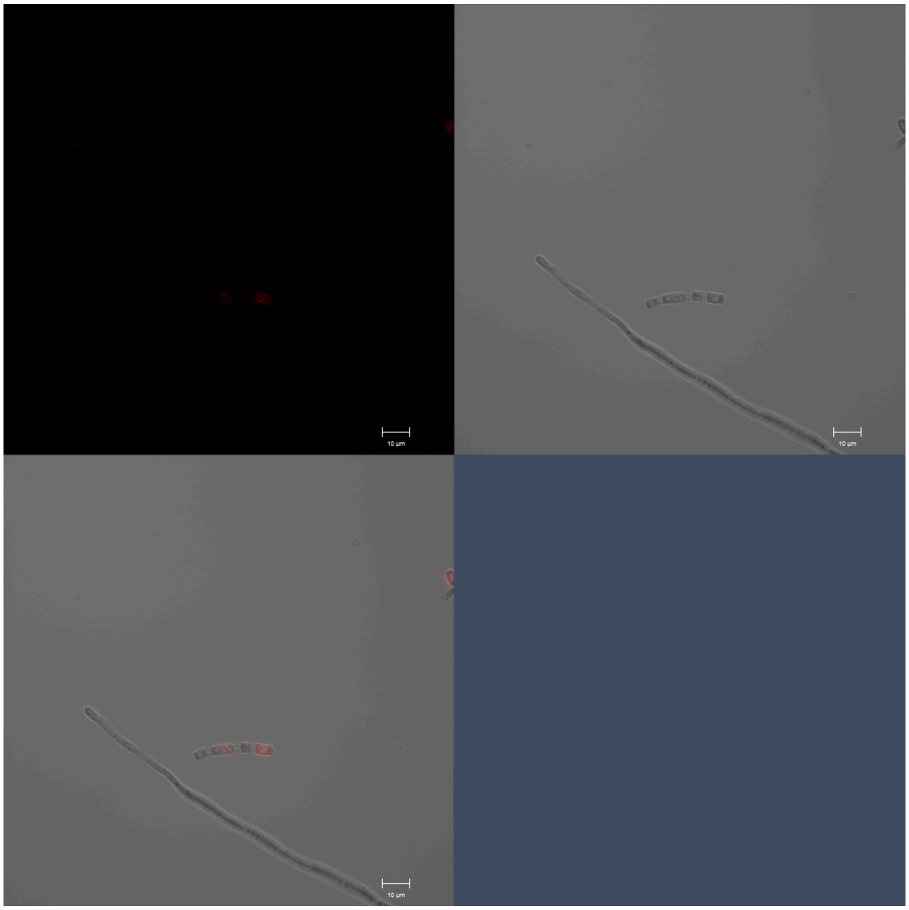

het-q1, Field 3, Replicate 2

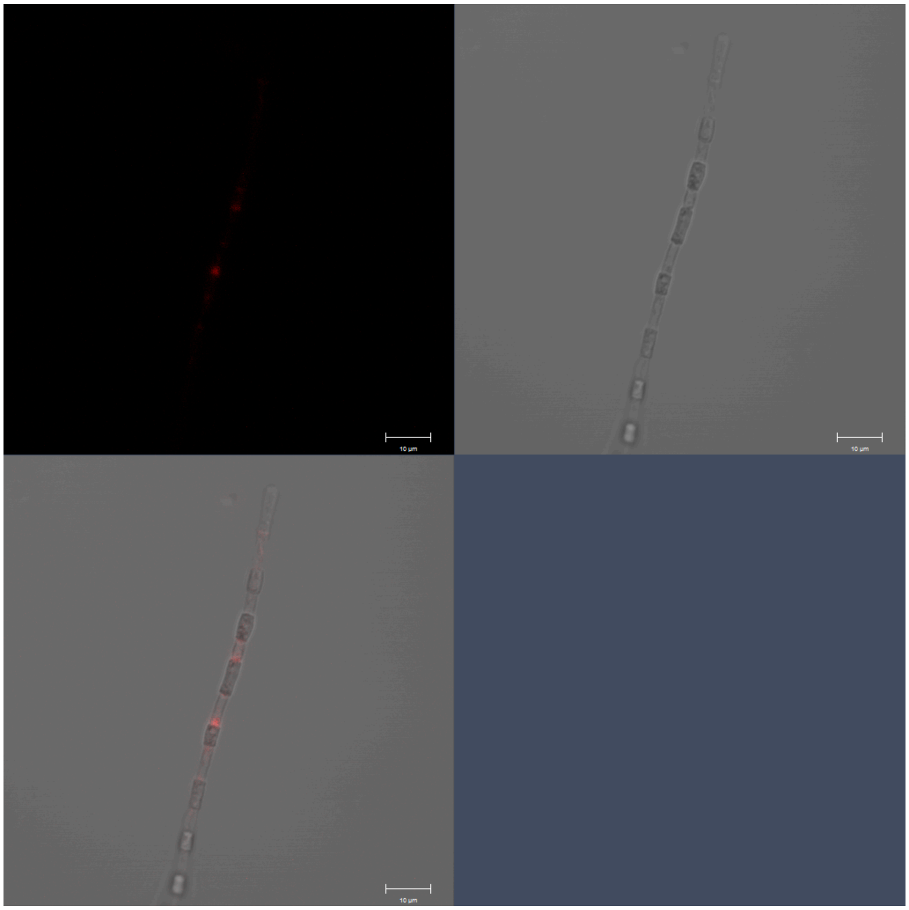

het-q1, Field 4, Replicate 2

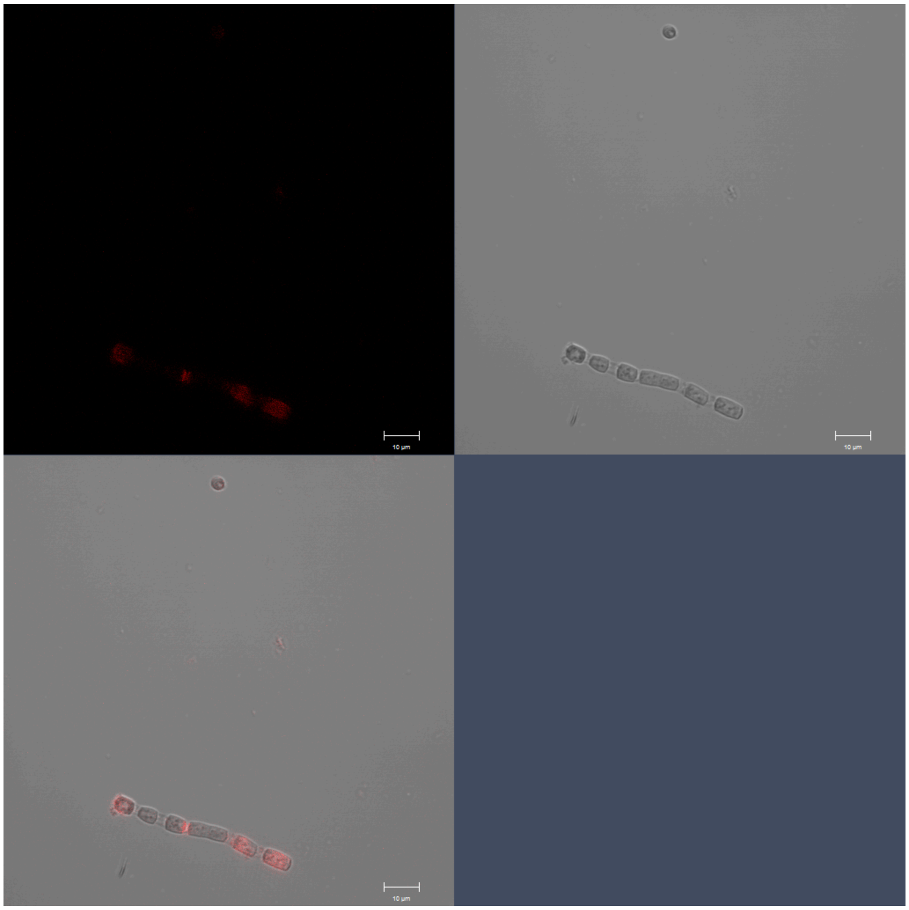

het-q1, Field 5, Replicate 2

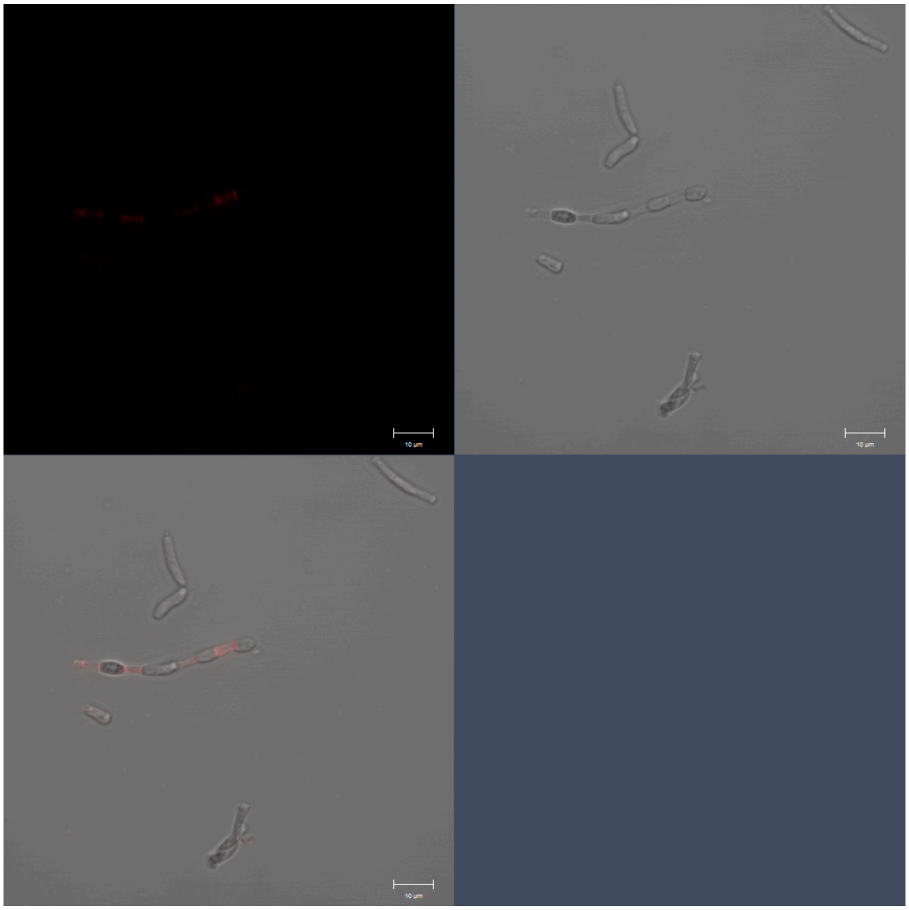

het-q2-like, Field 1, Replicate 1

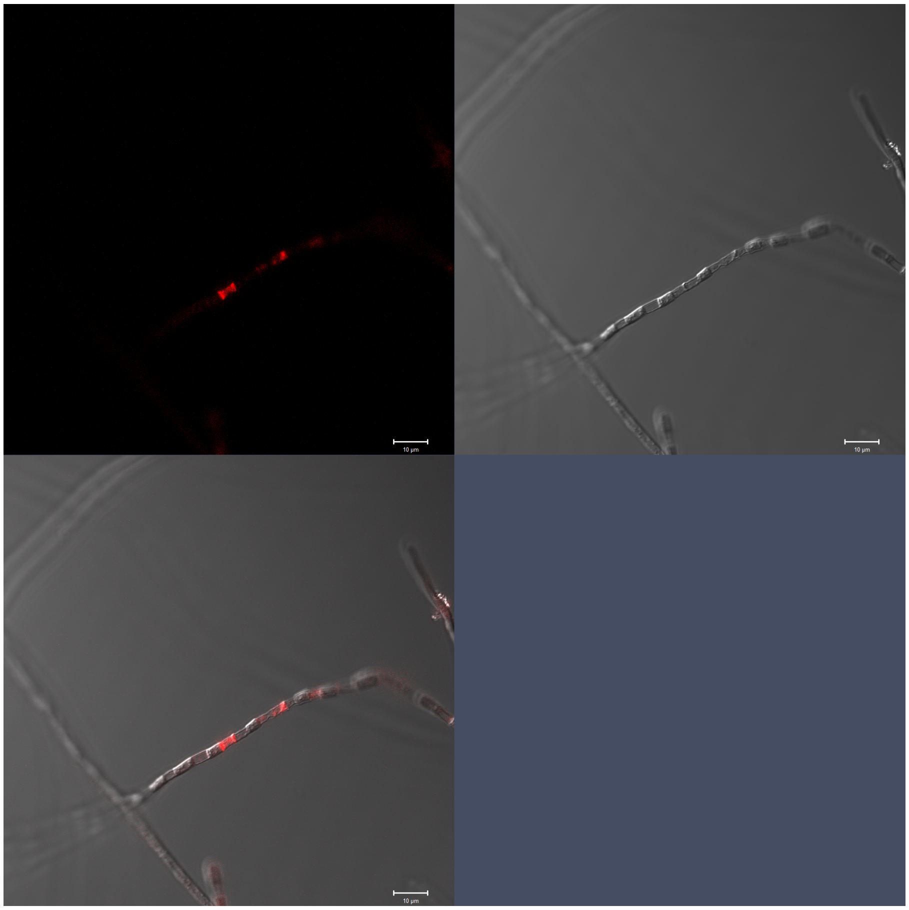

het-q2-like, Field 2, Replicate 1

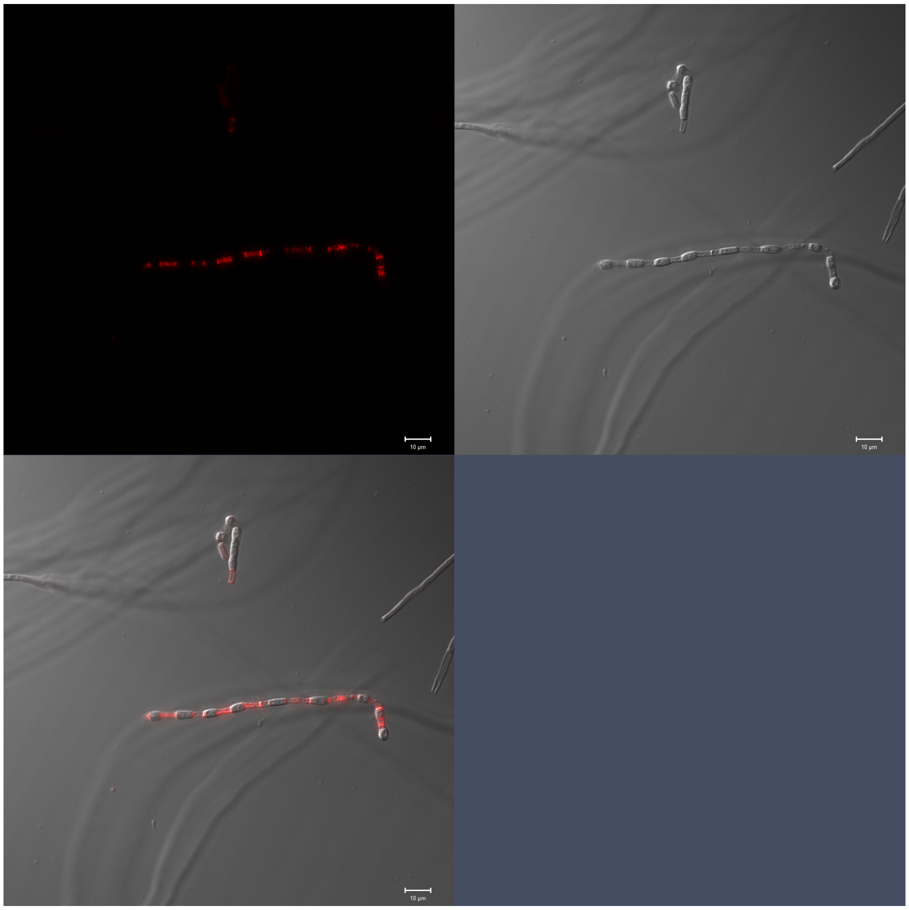

het-q2-like, Field 3, Replicate 1

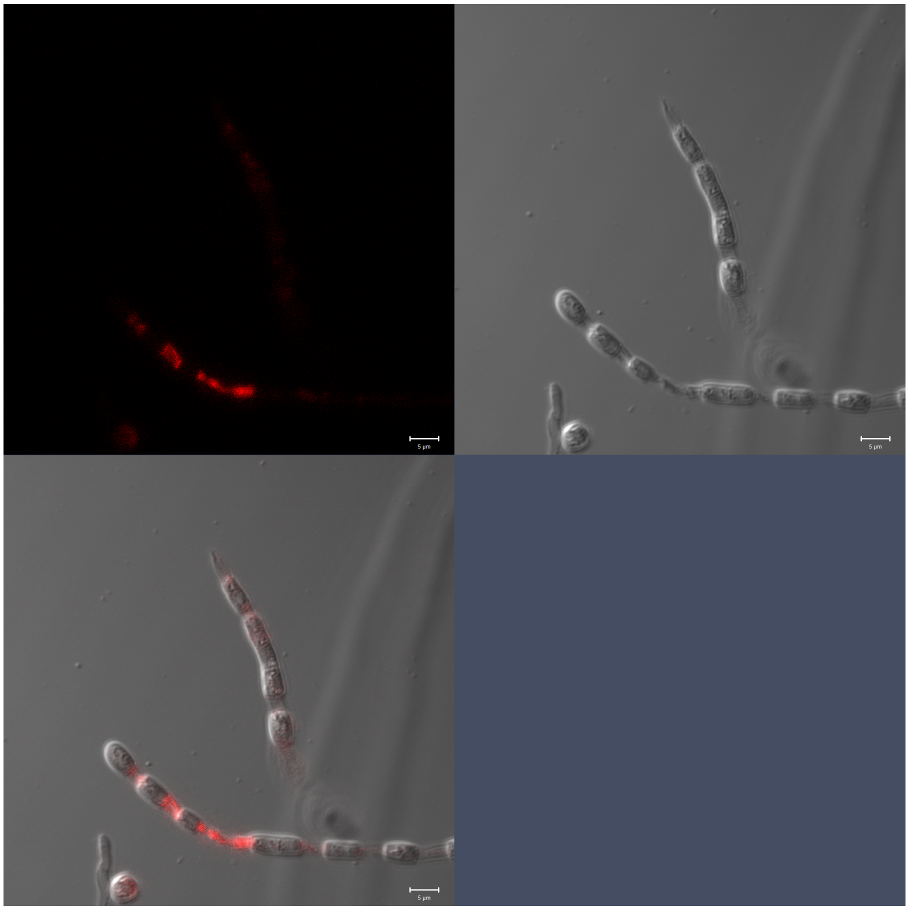

### het-q2-like, Field 4, Replicate 1

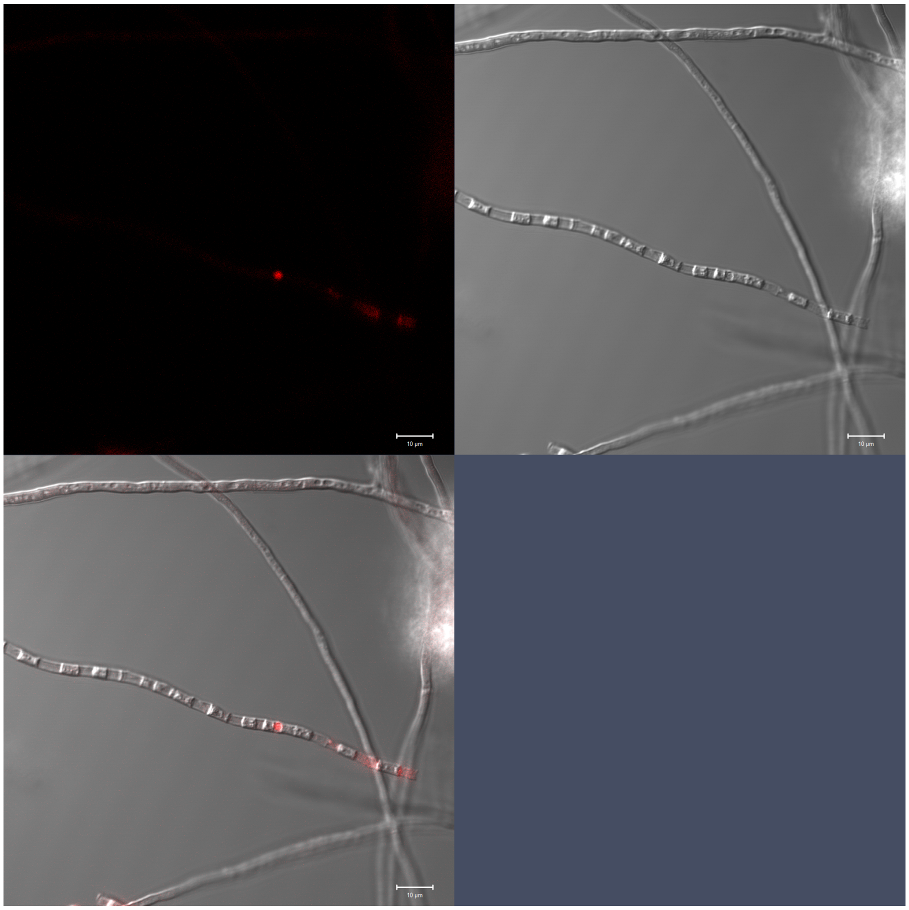

het-q2-like, Field 5, Replicate 1

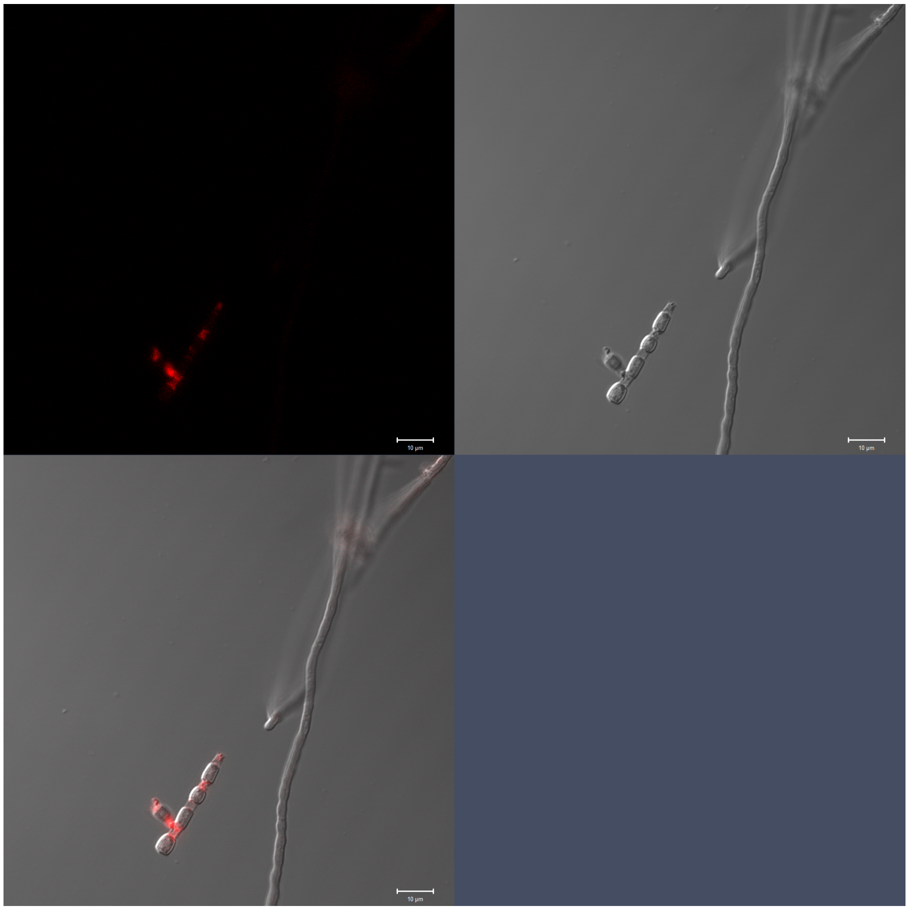

het-q2-like, Field 1, Replicate 2

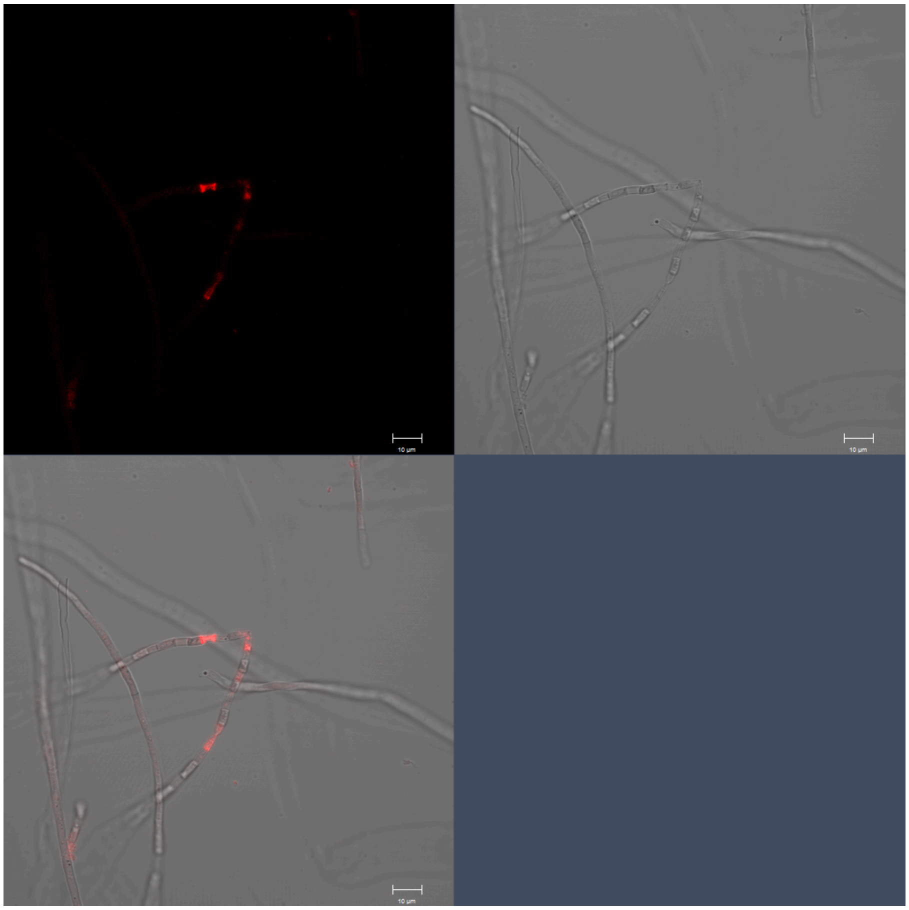

het-q2-like, Field 2, Replicate 2

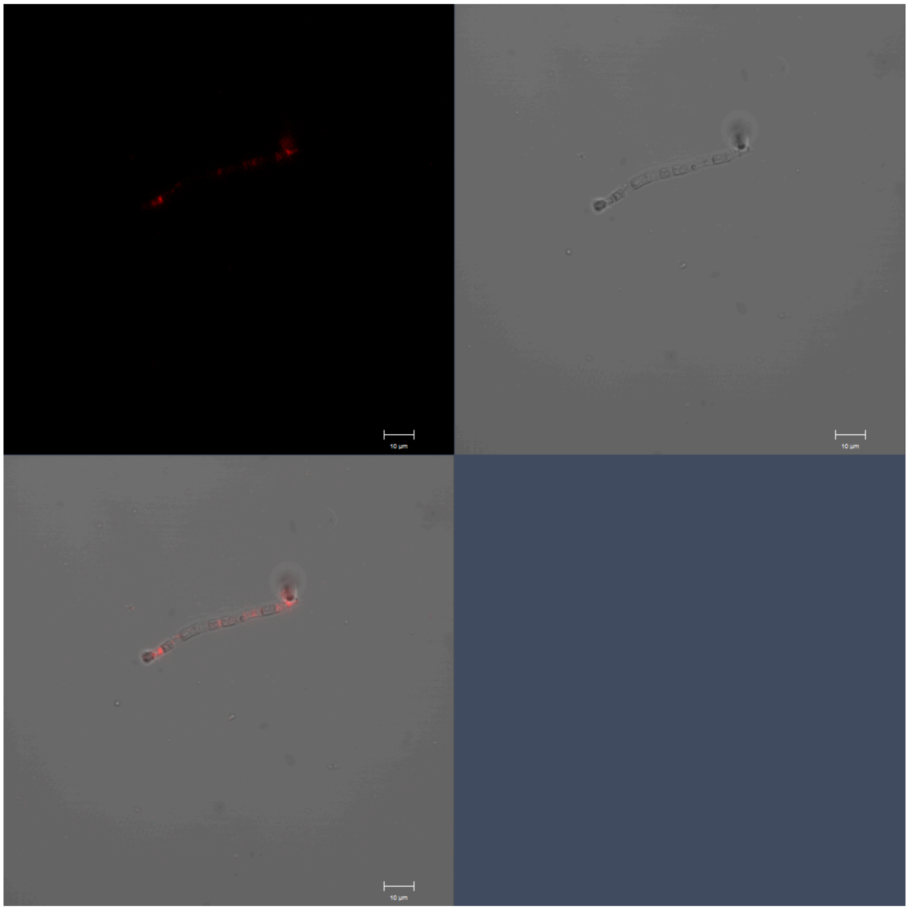

het-q2-like, Field 3, Replicate 2

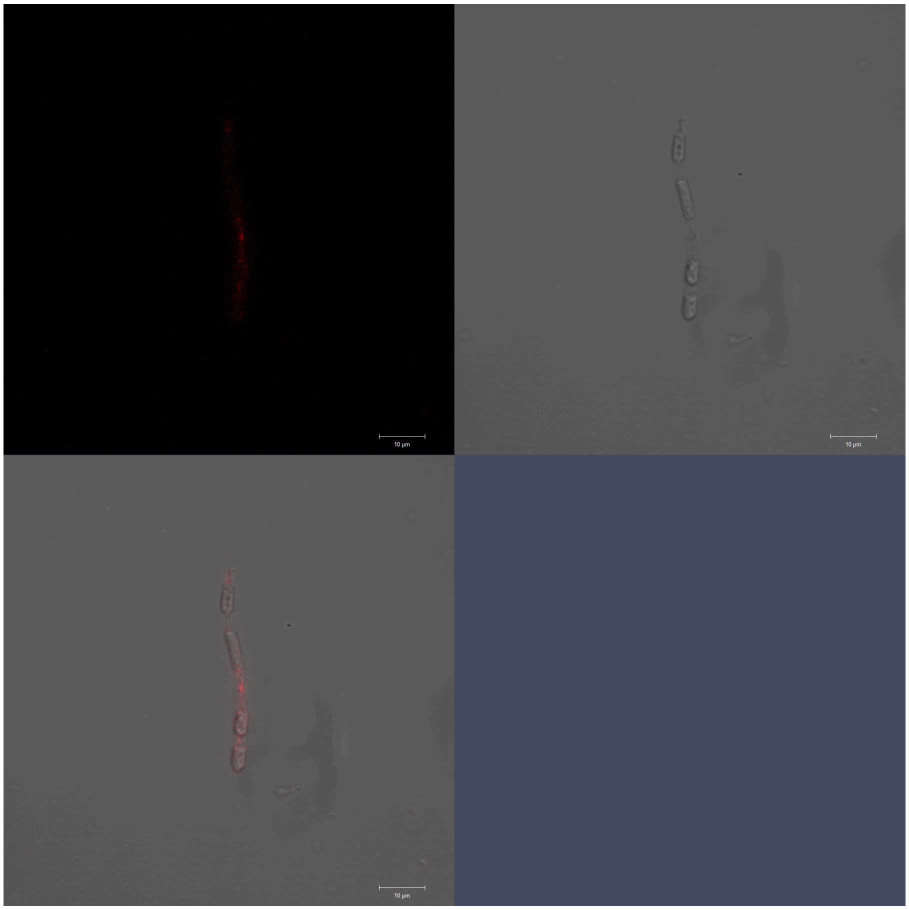

het-q2-like, Field 4, Replicate 2

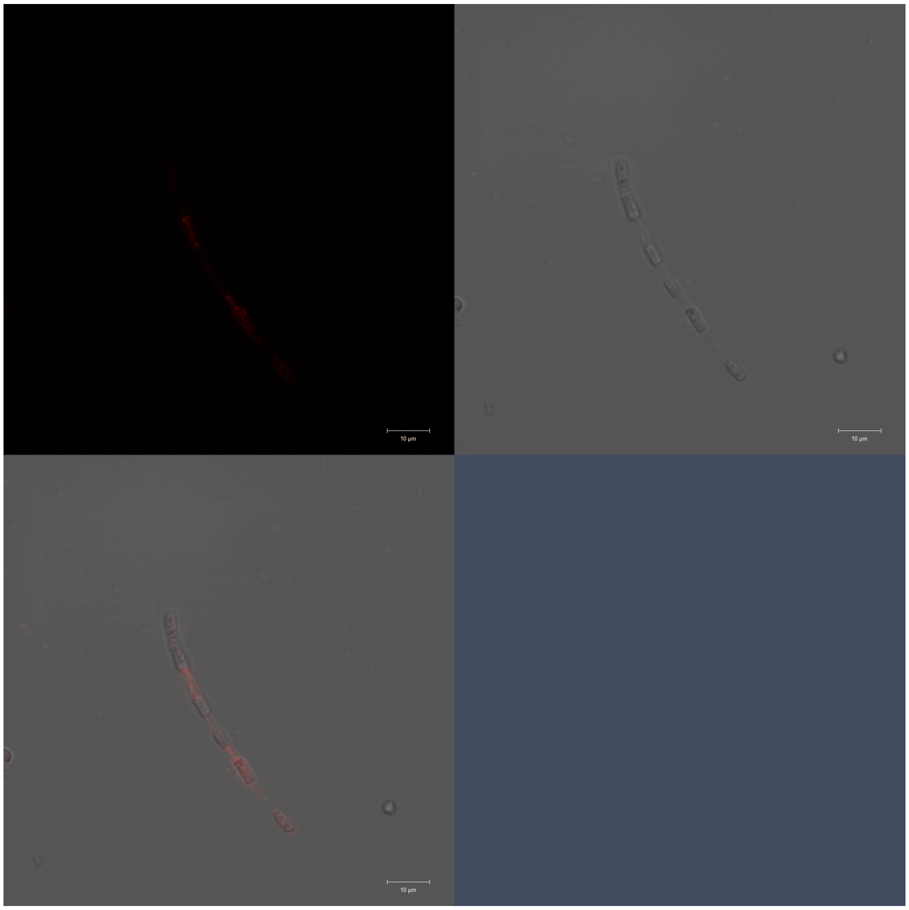

het-q2-like, Field 5, Replicate 2

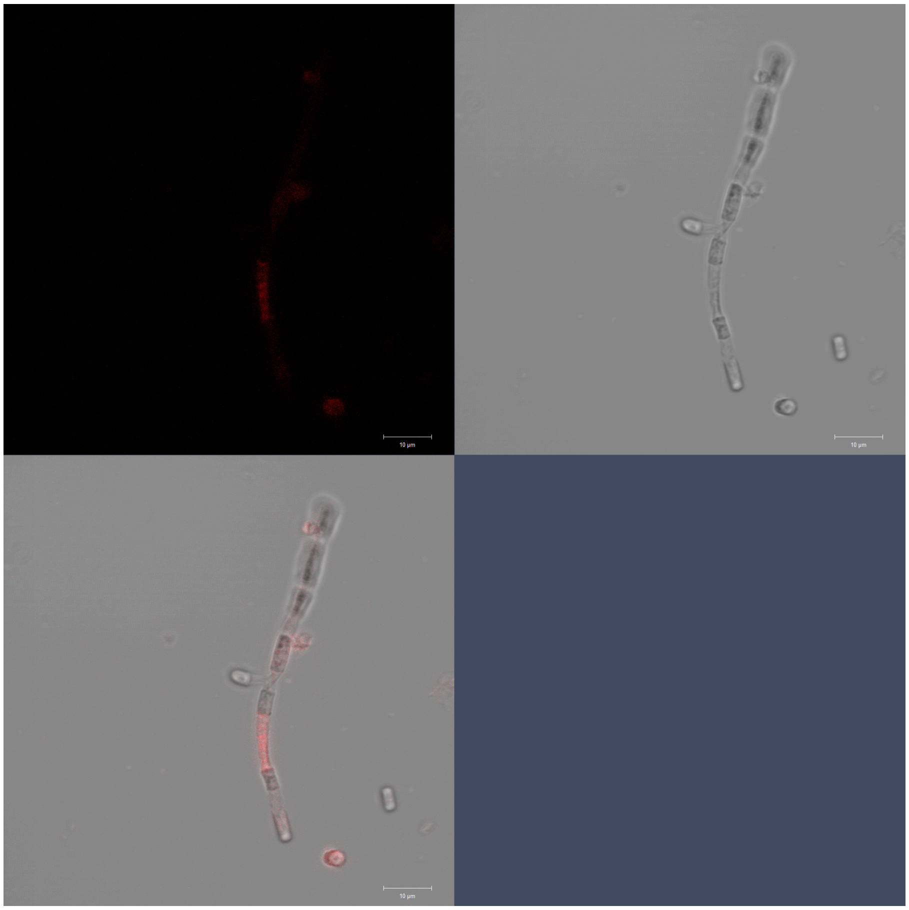
