## Supplementary figures and images for "Genetics and genomics of enteroarthroconidial development in *Coccidioides*"

### Fig S1

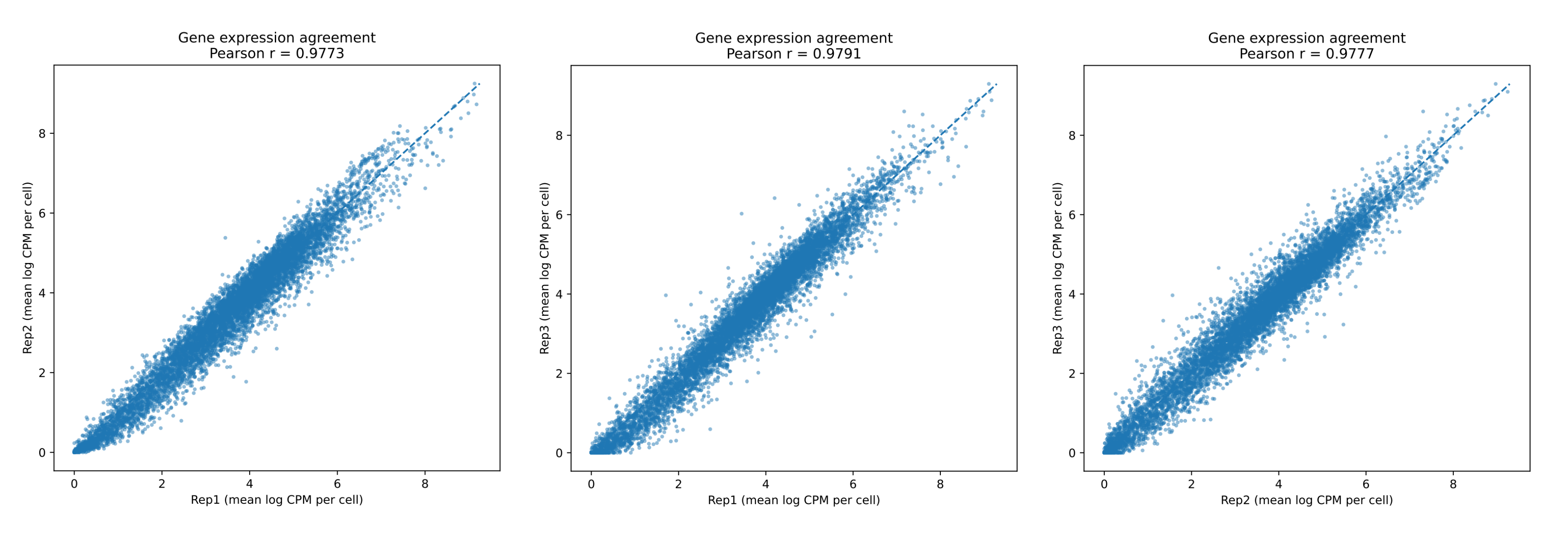

### S1 Movie

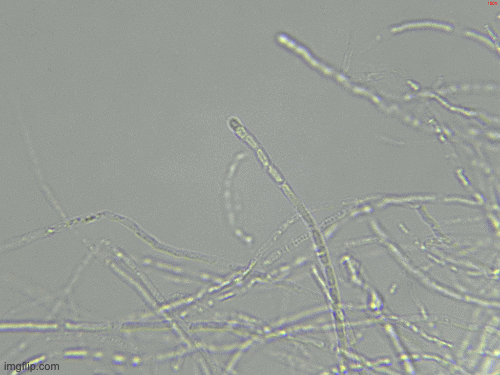
